# Chemically modified CRISPR-Cas9 enables targeting of individual G-quadruplex and i-motif structures, revealing ligand-dependent transcriptional perturbation

**DOI:** 10.1101/2024.10.14.618195

**Authors:** Sabrina Pia Nuccio, Enrico Cadoni, Roxani Nikoloudaki, Silvia Galli, An-Jie Ler, Claudia Sanchez-Cabanillas, Thomas E. Maher, Ella Fan, Dilek Guneri, Gem Flint, Minghui Zhu, Ling Sum Liu, Christopher R. Fullenkamp, Zoë Waller, Luca Magnani, John S. Schneekloth, Marco Di Antonio

## Abstract

The development of selective ligands to target DNA G-quadruplexes (G4s) and i-motifs (iMs) has revealed their relevance in transcriptional regulation. However, most of these ligands are unable to target individual G4s or iMs in the genome, severely limiting their scope. Herein, we describe a new Approach to Target Exact Nucleic Acid alternative structures (ATENA) that relies on the chemical conjugation of established G4 and iM ligands to a catalytically inactive Cas9 protein (dCas9), enabling their individual targeting in living cells. ATENA demonstrated that the selective targeting of the G4 present in the oncogene *c-MYC* leads to the suppression of transcripts regulated exclusively by one of its promoters (P1). Conversely, targeting the *c-MYC* iMs on the opposite strand leads to the selective increase of P1-driven transcripts. ATENA revealed that G4-mediated transcriptional responses are highly ligand-specific, with different ligands eliciting markedly different effects at the same G4-site. We further demonstrated that the basal expression levels of the gene targeted can be used to predict the transcriptional impact associated with G4-stabilization. Our study provides an innovative platform to investigate G4- and iM-biology with high precision and unveils the therapeutic relevance of individual DNA structures with unprecedented selectivity.

## Introduction

G-quadruplex (G4) structures can promptly form within single-stranded DNA sequences rich in guanines by generating stacks of G-quartets held together by Hoogsteen hydrogen-bonding that are further stabilized by coordination of potassium ions (Fig.1a) ^1^. The ability of G-rich sequences to form G4s under physiological conditions has been known for decades ^2,3^. Yet, the biological relevance of these structures has been heavily disputed until recently. Over the past decade, the development of orthogonal approaches to detect and map G4s has provided robust evidence to support their formation in living cells. These include immuno-fluorescence ^4^, live-cell imaging ^5,6^, and genome-wide mapping strategies ^7–10^.

**Figure 1:**
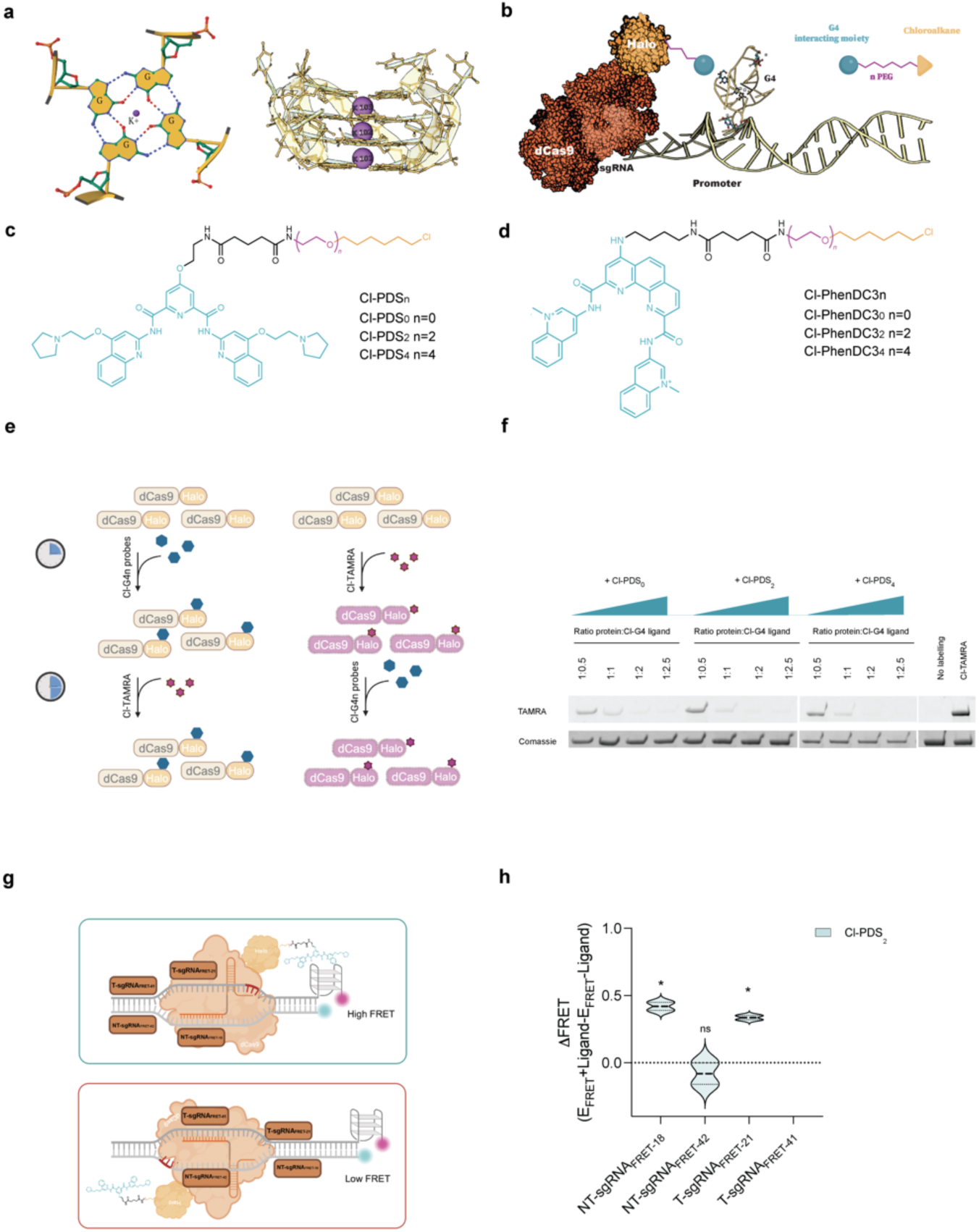
Chemical labeling of dCas9 for selective G4-targeting. **a**, Schematic representation of a G-Tetrad *via* Hoogsteen base pairing with central cation (K^+^) (modified from PDB file:6W9P) (Left) and G-Tetrad stacking to form G4-structure (modified from PDB file:6W9P using Protein Imager ^42^). (Right). **b**, Schematic overview of ATENA: dCas9-Halo fusion protein functionalized with chloroalkane-modified G4-ligands enables single G4-targeting through sgRNA guidance. **c,** Chemical structure of chloroalkane-modified PyPDS, where n indicates the different PEG linker lengths (Cl-PDS_n_). **d,** Chemical structure of chloroalkane-modified PhenDC3 (Cl-PhenDC3_n_), where n indicates the different PEG linker lengths (Cl-PhenDC3_n_). **e,** Illustration of the competition assay workflow to test the binding ability of Cl-PDS_n_ probes to dCas9-Halo purified recombinant protein. **f,** SDS-Page gel of the Cl-PDS_n_ competition assay shows each sample’s fluorescent level acquired in the TAMRA channel (542 nm) and the corresponding protein level (Coomassie staining): (n=2). **g,** Schematic representation of the dually labeled FRET oligos containing c-KIT2-G4 forming sequence bound by dCas9-Halo labeled with Cl-PDS_n_ probes to study G4 stabilization with respect to the PEG-linker length and sgRNA positioning. **h, Δ**FRET efficiency of the decorated dCas9-PDS (with Cl-PDS_2_) complex targeting c-KIT2-G4. The values indicated were extrapolated from the band intensity measured in the Cy3 and Cy5 channels (Typhoon FLA 9500). The signals in both channels were normalized for the background and the sgRNA NTC control. These normalized fluorescence values were then used to calculate the ΔFRET efficiency for each sgRNA: FRET-Efficiency (E)+ligand _sgRNAx_ – FRET-Efficiency (E)-ligand _sgRNAx_ (n=2). Data presented are the mean of n = number of independent experiments. Statistical significance was calculated using a two-tailed t-test in GraphPad Prism; p-value: ns > 0.05, * ≤0.05, ** ≤0.01, *** ≤0.001, **** ≤0.0001.

While G4-formation in cells has been extensively validated, its active contribution in regulating biological processes is yet to be fully demonstrated. Given the high enrichment of G4s at gene promoters measured experimentally *in vitro* and in cells ^9,11^, it has long been speculated that these structures may play an essential role in regulating gene expression ^12^. Indeed, many G4-selective ligands have been developed to date, revealing that targeting G4s within gene promoters is generally associated with transcriptional suppression ^13^. Since key oncogenes like *c-MYC, KRAS,* and *BCL-2* bear a G4 in their promoter, applying G4 ligands for cancer intervention has been extensively investigated ^14^. However, the human genome displays more than 700,000 experimentally detected G4-structures *in vitro* ^11,15^ and around ∼10,000 G4s detected in chromatin using genomics strategies ^7–10,16^, making single-G4 targeting by small molecules extremely challenging. Indeed, most ligands recognize G4s by establishing π-π end-stacking with G-tetrads, a promiscuous interaction that hampers these molecules from displaying any significant inter-G4 selectivity^17^. For example, evidence supports that the transcriptional suppression of *c-MYC* observed upon treatment with certain G4-ligands is an indirect consequence of global G4 stabilization rather than a specific response regulated exclusively by the G4 in the *c-MYC* promoter ^18^.

Additionally, there are experimental discrepancies between what could be the endogenous biological function of G4s and what is instead observed when ligands bind these structures. A series of independent genomic studies consistently indicated that G4-formation is strongly associated with active gene expression ^12,19^, contrasting the transcriptional repression mostly observed upon G4-ligand treatment. Similarly, recent studies have leveraged gene-editing techniques to demonstrate how the selective deletion of the G-rich sequence in the *c-MYC* promoter responsible for the G4-structure formation on this promoter (MYC-G4) is associated with loss of transcriptional activity ^20^, further pointing to an active role of G4s in stimulating transcription. To justify the discrepancies between endogenous G4-function versus the responses observed by G4-targeting with small-molecules, it has been postulated that ligand binding can prevent key transcriptional factors from recognizing G4s, leading to transcriptional repression^21^. However, it remains challenging to discern whether transcriptional suppression at specific genes is caused by protein displacement at a given G4 site or is a response triggered by global G4-stabilization, as reported for *c-MYC*^18^. It is also often supposed that structurally different G4-ligands should elicit similar responses when targeting the same G4s, with the assumption that G4 binding proteins respond identically to a ligand-bound G4 irrespective of the ligand used. Overall, this highlights the urgent need for tools that can provide inter-G4 selectivity to widely used G4 ligands to fully underpin G4-biology and harness the therapeutic potential of these DNA secondary structures.

Similarly, i-motifs (iMs) structures have been recently mapped through the human genome^22^, visualized in cells^23^, and have been heavily linked with transcriptional regulation^22^. Although several ligands have been shown to bind the i-motif-forming sequences in *c-MYC*^23^, the exact biological responses elicited by targeting iMs still remain elusive, owing to their lack of inter-iMs selectivity that confounds phenotypes arising from global versus site-specific iM targeting.

Therefore, it is evident that the absence of inter-G4 or inter-iMs selectivity displayed by most of the available ligands strongly hampers their use as therapeutic agents and their application as reliable tools to investigate the biology of these DNA structures. Hence, the development of ligands displaying preferential binding toward a specific G4 or iM structure (i.e., *c-MYC*) has represented a longstanding quest in this area of research. To this end, Schneekloth and co-workers recently designed a small-molecule ligand (DC-34) that displays preferential binding to the G4 in the *c-MYC* promoter^24^. Treatment of multiple myeloma cell lines with DC-34 revealed suppression of *c-MYC* transcription with minimal perturbation of other G4-bearing oncogenes, such as *KRAS* and *BCL-2*^24^. Similarly, our group and others have shown that short peptides and oligonucleotides can also display binding selectivity towards specific G4s^25–28^. Nevertheless, these remain isolated examples that require ligand optimization to achieve binding selectivity against any given DNA structure of interest and are not suitable for high-throughput screening of G4 or i-motif function at scale. Furthermore, the highly diverse chemical and structural nature of ligands targeting G4s and iMs might represent an additional confounding factor. For example, different G4-ligands might elicit different biological responses when binding to the same G4s, leading to experimental observations that are dependent on the type of ligand used, rather than reflecting the endogenous biological function of the targeted G4.

To overcome these limitations, we have developed ATENA (Approach to Target Exact Nucleic Acid alternative structures), a CRISPR-Cas9-based platform that enables selective localization of different G4 or iM ligands in the proximity of a given DNA structure of interest (Fig.1b). To achieve this, we have exploited HaloTag technologies to selectively install modified G4 or iM binding ligands onto Cas9 protein in living cells^29^. More specifically, we have designed a small library of established G4-ligands (pyridostatin (PDS)^30^ and PhenDC3^31^) functionalized with chloroalkane side chains that ensure incorporation into a nuclease-inactive Cas9 protein fused to a HaloTag (dCas9-Halo) ^32^.

Similarly, we decorated one of the iM selective peptides (Pep-RVS) ^33^, recently developed by the Waller’s group, with a chloroalkane side chain to explore selective iM targeting with ATENA. The synthesized analogs of PDS, PhenDC3, and Pep-RVS contain different polyethylene-glycol (PEG)-linkers (n = 0, 2, 4, Fig.1 c,d), with the PEG chain serving to tether the DNA-binding scaffold to the HaloTag-binding chloroalkane. By fine-tuning the PEG linker length, we optimized the spacing between the DNA-binding moiety and the dCas9-Halo fusion to attain ideal G4 engagement with ATENA, using a dedicated FRET-based assay. We leveraged this knowledge to deploy ATENA in cells and achieve selective targeting of either the G4 or the iM present in the promoter of the proto-oncogene *c-MYC*. ATENA-mediated G4-targeting resulted in the reduction of *c-MYC* transcripts generated exclusively by one of the four promoters regulating *c-MYC* expression (P1), irrespective of the ligand used, which is in agreement with recent literature ^20^. Additionally, ATENA has shown, for the first time, that selective MYC-G4 targeting does not result in a net reduction of *c-MYC* expression, as increased transcription from the alternative P2 promoter counterbalances the P1-specific downregulation induced by G4 engagement. The P1-dependent downregulation upon G4-targeting was also confirmed by treatment with the MYC-G4 selective ligand DC-34, further validating the ability of ATENA to target individual G4s. Importantly, we have also observed that guiding G4-ligands in the proximity of the *c-MYC* promoter TATA-box can lead to G4-independent transcriptional suppression. This strongly suggests that previous observations obtained with dCas9 decorated with multiple G4-ligands are likely reflecting occupancy of transcriptional regulatory regions rather than genuine G4-binding ^34^.

Upon employing ATENA to target selectively the iM present in the promoter of *c-MYC*, using a HaloTag-compatible version of Pep-RVS, we have also observed P1-specific transcriptional perturbation. An increase in P1-expression was associated with iM targeting, opposing the transcriptional inhibition observed by G4 stabilisation, in line with other systems which indicate that iM-binding results in induction of gene expression^35,36^ and that iMs and G4 shape opposing effects in cells^37^.

While downregulation of P1-mediated *c-MYC* expression was observed with two distinct ligands, PDS and PhenDC3, ATENA also revealed that other G4s could provide different transcriptional outcomes when targeted by the same compounds. This suggested that the biological response elicited by G4 stabilization might reflect more the structural nature of the G4-ligand complex rather than providing direct insights into the endogenous function of G4s, underscoring the importance of the choice of ligand used in the different reports aimed at unveiling G4-biology. Finally, we showcased the ability of ATENA to infer the biological relevance of cell line-specific G4s, revealing a strong dependence of the transcriptional perturbation attained upon ligand treatment on the expression level of the gene targeted.

Altogether, our study provides robust evidence supporting ligand and transcriptional-dependent responses to both G4s and iM targeting. ATENA functions as a modular platform to target individual DNA secondary structures in living cells, enabling a precise study of G4- and iM-biology. We anticipate that ATENA will offer unprecedented potential for screening cell and ligand-specific responses to DNA secondary structure targeting in a high-throughput manner, which can be further translated for therapeutic design and development.

## Results

### Chemical labelling of CRISPR-Cas9 proteins with G4 ligands

Catalytically inactive CRISPR-Cas9 (dCas9) fused to functional proteins has been widely used in biology to attain site-selective perturbation of gene expression ^38^. This strategy takes advantage of the selectivity provided by dCas9 bound to a short-guiding RNA (sgRNA) in recognizing a specific genomic site by base-pairing, which can be used to recruit an effector protein (i.e., a transcription factor or an epigenetic enzyme) at the targeted site ^39^. A similar strategy has been recently devised to decorate dCas9 with G4-ligands using non-covalent biotin-streptavidin recognition^34^. Irreversible chemical functionalization of dCas9 proteins has been previously achieved using commercially available chloroalkane-modified fluorophores to label a dCas9-Halo fusion protein in living cells ^39^. Therefore, we hypothesized that generating modified G4-ligands functionalized with chloroalkane moieties could have been exploited to decorate with higher control and irreversible covalent chemistry Cas9 proteins with G4 ligands under physiological conditions. To achieve this, we designed analogues of the widely established G4 ligands PDS based on the previously described PyPDS scaffold ^5^. Unlike PDS, PyPDS presents a single primary amine within its structure, which can be selectively functionalized with chloroalkane sidechains by peptide coupling (Fig. 1c) ^5^. After successful synthesis of PyPDS following previously established methods^5^, we functionalized the primary amine of the molecule with chloroalkane side chains of different lengths, enabling systematic investigation of the ideal distance between the G4-binding scaffold PyPDS and the HaloTag protein to achieve optimal G4-engagement. Specifically, we have used linkers containing different PEG repeats (n = 0, 2, 4) to vary the distance between PyPDS and the chloroalkane (Cl-PDS_n_, Fig. 1c, SI-1).

To avoid limiting the use of ATENA to a single G4-ligand, we also functionalized with chloroalkane moieties another widely characterized G4 ligand called PhenDC3 using the same chemical strategy (Cl-PhenDC3_n_) ^31^. Unlike PDS, PhenDC3 has a cationic side chain and is structurally bulkier, displaying a phenanthroline core wider than the pyridine one present in the PDS scaffold (Fig. 1c, d). While PDS and PhenDC3 have been extensively validated as selective G4-ligands, the structural differences between these two ligands might lead to distinct biological responses when targeting G4s in cells, which cannot be assessed with current methods. Therefore, we decided to systematically compare these two ligands with ATENA when recruited to a single-G4 site. To achieve this, we have synthesized a previously reported PhenDC3 analogue displaying an exocyclic primary amine ^40^, which can be functionalized by peptide coupling using the same synthetic strategy described for PyPDS to afford chloroalkane-modified PhenDC3 analogs that are compatible with HaloTag conjugation (Fig. 1d, SI-1).

### *In vitro* validation of covalent conjugation of G4-ligands to dCas9-Halo

With both PyPDS and PhenDC3 analogues in hand, we initially assessed whether functionalization with the chloroalkane side chains could affect the G4-binding properties of these molecules. To test this, we have subjected all the ligands to FRET or Circular Dichroism (CD) melting to evaluate their ability to stabilize G4-structures after chloroalkane functionalization. All analogues tested displayed good G4-stabilization, providing an increase in melting temperature (ΔT_m_ >10 °C at 4 μM) against four distinct G4 structures tested (c-MYC, hTelo, BCL-2 and c-KIT2; Extended Data Tables 1, 2, 3 and 4), with ΔT_m_ values comparable to the unfunctionalized ligands. This suggested that the addition of the chloroalkane side chains had a negligible effect on the G4-stabilization properties of both PhenDC3 and PyPDS.

Having confirmed that both chloroalkane-functionalized PyPDS and PhenDC3 analogues retained good G4-binding recognition properties, we next investigated if these molecules could be covalently engaged to dCas9-Halo and performed a competition assay *in vitro*. To this end, we expressed and purified the dCas9-Halo protein (see methods) and incubated it for 45 minutes with increasing concentrations of Cl-PDS_n_ and Cl-PhenDC3_n_. This was followed by incubation with an excess (5 μM) of the commercially available HaloTag^®^TAMRA (Cl-TAMRA) ligand to label dCas9-Halo with the TAMRA fluorophore. Since labeling of HaloTag is a covalent irreversible process ^29^, we reasoned that initial exposure of the dCas9-Halo protein to the chloroalkane functionalized G4-ligands would prevent subsequent incorporation of the fluorescent Cl-TAMRA, leading to a dose-dependent reduction of TAMRA incorporation (Fig. 1e). Indeed, all tested analogues induced a robust dose-dependent decrease of the dCas9-Halo TAMRA signal (Fig. 1f and Extended Data Fig. 1a), indicating high efficiency in labeling dCas9-Halo irrespectively of the G4-ligand (i.e., Cl-PDS_n_ or Cl-PhenDC3<u>_n_</u>) or the PEG-linker used to connect the chloroalkane to the G4-binding scaffold. To quantify labeling efficiency, we measured fluorescence intensity for each line of the gel and generated dose–response curves to extract the concentration of ligand required to attain 50% labeling of the dCas9-Halo *in vitro* (IV-CP_50_). As depicted in Extended Data Fig. 1b, c, all the analogues displayed very low IV-CP_50_ values (≤ 2 μM), demonstrating a strong ability to functionalize dCas9-Halo *in vitro*. This data supports the use of chloroalkane-modified G4 ligands to covalently label dCas9-Halo.

### *In vitro* optimization of sgRNAs and PEG-linker to attain G4-engagement

We next asked if dCas9-Halo functionalized with G4-ligands could be used to target individual G4s. To achieve this, we investigated the ideal distance between the dCas9 binding site and the targeted G4 to achieve optimal engagement of the G4 ligands with the targeted structure by systematically varying both the short guiding RNA (sgRNA) sequences used and the PEG-linker connecting the Halo reactive moiety (chloroalkane) to the G4-binding scaffold tested. To quantify G4-engagement, we designed a dually fluorescent-labelled DNA template containing an established G4-forming sequence (c-KIT2-G4) at its 3’ end to monitor G4-stabilization through FRET (Fig. 1g). We then designed two sgRNAs to orient the dCas9-Halo complex towards the G4 structure sitting at either 18 (NT-sgRNA_FRET-18_) or 42 (NT-sgRNA_FRET-42_) base pairs from the targeted G4 and whose Protospacer Adjacent Motifs (PAM) are located on the non-template strand bearing the G4 (NT, Fig. 1g). This is based on previous structural studies indicating that the C-terminal domain of the Cas9 protein, where the Halo protein is situated, will point towards the 3’ end of the PAM sequence ^41^. Moreover, we have used a standard non-targeting RNA control (NTC) to rule out any potential unspecific binding that is not strictly mediated by sgRNA-driven proximity. We then used Cl-PDS_n_ molecules as prototype G4-ligand to assess the extent of G4 targeting by ATENA under different conditions, by measuring changes in FRET when targeting the oligo construct with the dCas9-Halo complex in the presence or the absence of Cl-PDS_n_. We failed to detect any significant changes (p>0.05) in the FRET signal for both sgRNAs tested (NT-sgRNA_FRET-18_ vs NT-sgRNA_FRET-42_) when using Cl-PDS_0_ (Extended Data Fig. 1d). Considering that Cl-PDS_0_ can efficiently bind to dCas9-Halo (Fig. 1f, IV-CP_50_ = 1.6 µM), the lack of G4-stabilization displayed by this molecule indicates that the linker connecting the PDS-scaffold to the dCas9-Halo is inadequately short for engaging with the G4-structure. Nevertheless, a trend showing higher changes in FRET efficiency when using sgRNAs closer to the G4 (NT-sgRNA_FRET-18_ vs NT-sgRNA_FRET-42_) suggests that placing the dCas9 closer to the G4 facilitates ligand engagement (Extended data Fig. 1d). Indeed, when decorating dCas9-Halo with a PDS analogue with a longer PEG-linker (Cl-PDS_2_), a significant increase (p<0.05) in FRET-signal could be measured when using NT-sgRNA_FRET-18_ (ΔFRET = 0.42), which is indicative of G4-engagement. However, when using NT-sgRNA_FRET-42_ we failed to measure an increase in FRET signal, confirming that placing dCas9-PDS closer to the G4 provides better ligand engagement. To further investigate ideal conditions to obtain G4-targeting with ATENA, we also designed T-sgRNA_FRET-21_ and T-sgRNA_FRET-41_ that sit at 21 and 41 base pairs from the G4 but whose PAM is located on the template strand (T, Fig. 1g) to investigate the effect of the dCas9-Halo orientation on G4 targeting (Fig. 1g, h). Consistent with our observations indicating that closer placement of dCas9-Halo to the G4 is linked with better ligand engagement, we detected a significant (p<0.05) increase in FRET signal when using T-sgRNA_FRET-21_ (ΔFRET = 0.33, Fig. 1h) that was abrogated when using T-sgRNA_FRET-41_ (Fig. 1h, SI). This further indicated that optimal G4-targeting by ATENA is achieved by using sgRNAs closer to the G4 regardless of the orientation imposed by the sgRNAs used (Fig.1h and SI Fig. 1g). When further increasing the PEG-linker using Cl-PDS_4_, we failed to observe any significant (p<0.05) increase in FRET efficiency with both NT-sgRNA_FRET-18_ and NT-sgRNA_FRET-42_, suggesting that using PEG-linkers that are excessively long is detrimental to G4-engagement, possibly due to high entropic penalty associated with ligand-recognition (Extended Data Fig. 1e). Altogether, our study demonstrated that dCas9-driven G4-ligand engagement is both PEG-linker and sgRNA dependent, with optimal G4-engagement achieved when using a PEG2 linker and sgRNAs placing the dCas9-Halo complex as close as possible (depending on the PAM availability) to the targeted G4.

### Chloroalkane-modified ligands can label dCas9-Halo efficiently in cells

Having optimized conditions to achieve G4-engagement *in vitro* with ATENA - using Cl-PDS_n_ and c-KIT2 G4s as a model system - we next investigated whether ATENA can be used to stabilize individual G4 structures in living cells. The use of cell lines that constitutively express dCas9-Halo is essential to ensure consistent cellular levels of the protein across different experiments, avoiding bias introduced by the significantly variable levels of protein expression typically with transient transfection used in previous reports. ^34^ To this end, we integrated dCas9-Halo into the genome of the breast cancer cell line (MCF7) using standard lentiviral integration approaches (see methods). We selected MCF7 cells in light of the highly diverse transcriptional response previously reported upon treatment with PDS ^43^, which we wanted to investigate further with ATENA. Successful integration of dCas9-Halo was confirmed by PCR-based genotyping and Western Blot (Extended Data Fig. 2a, b). Next, we evaluated the ability of the chloroalkane functionalized ligands to bind dCas9-Halo in cells. Using an established chloroalkane penetration assay (CAPA), we compared each ligand’s relative potency to label covalently dCas9-Halo under physiological conditions ^44^. During CAPA, cells are initially exposed to increasing concentrations of the chloroalkane-modified G4-ligands, before incubation with a Halo-reactive Oregon Green fluorophore (Cl-OG), which reacts with any Halo-tag binding site that has been left unoccupied by the previous exposure to the chloroalkane G4-ligands (Fig. 2a, i-ii). The efficiency of G4-ligand incorporation can be, therefore, measured as an inverse function of the Oregon Green fluorescence emission, as successful G4-ligand incorporation to Halo prevents subsequent fluorophore functionalization (Fig. 2a, iii-iv). To quantify this numerically, we calculated the half-maximal chloroalkane penetration value (CP_50_), which is the ligand concentration required to label 50% of the available dCas9–Halo molecules and can be used as a direct readout of target occupancy^44^. As displayed in Fig. 2a (right), treatment of MCF7 cells expressing dCas9-Halo with both Cl-PDS_0_ and Cl-PDS_4_ revealed a modest dose-dependent reduction of Oregon Green emission, providing CP_50_ values of 15.9 μM and 5.4 μM, respectively (Extended Data Fig. 2c). Conversely, Cl-PDS_2_ could label ∼90% dCas9-Halo at a concentration as low as 0.25 μM (CP_50_ 0.012 μM, Extended Data Fig. 2c), saturating at 2.5 μM (Fig. 2a), suggesting that the cellular permeability and bioavailability of Cl-PDS_2_ were particularly suitable for its application in ATENA. Given that the PEG2 linker also led to best G4-engagement *in vitro* (Fig. 1h), we decided to assess the compatibility with ATENA of a different G4-ligand (PhenDC3) bearing a PEG2 linker (Cl-PhenDC3_2_) through CAPA. Gratifyingly, Cl-PhenDC3_2_ labelled very efficiently dCas9-Halo in cells, yielding a CP_50_ value of 1.7 μM (Extended Data Fig. 2d). Cl-PhenDC3_4_ showed a similar trend to Cl-PDS_4_, indicating that PEG4 functionalized ligands could not be employed in ATENA.

**Figure 2:**
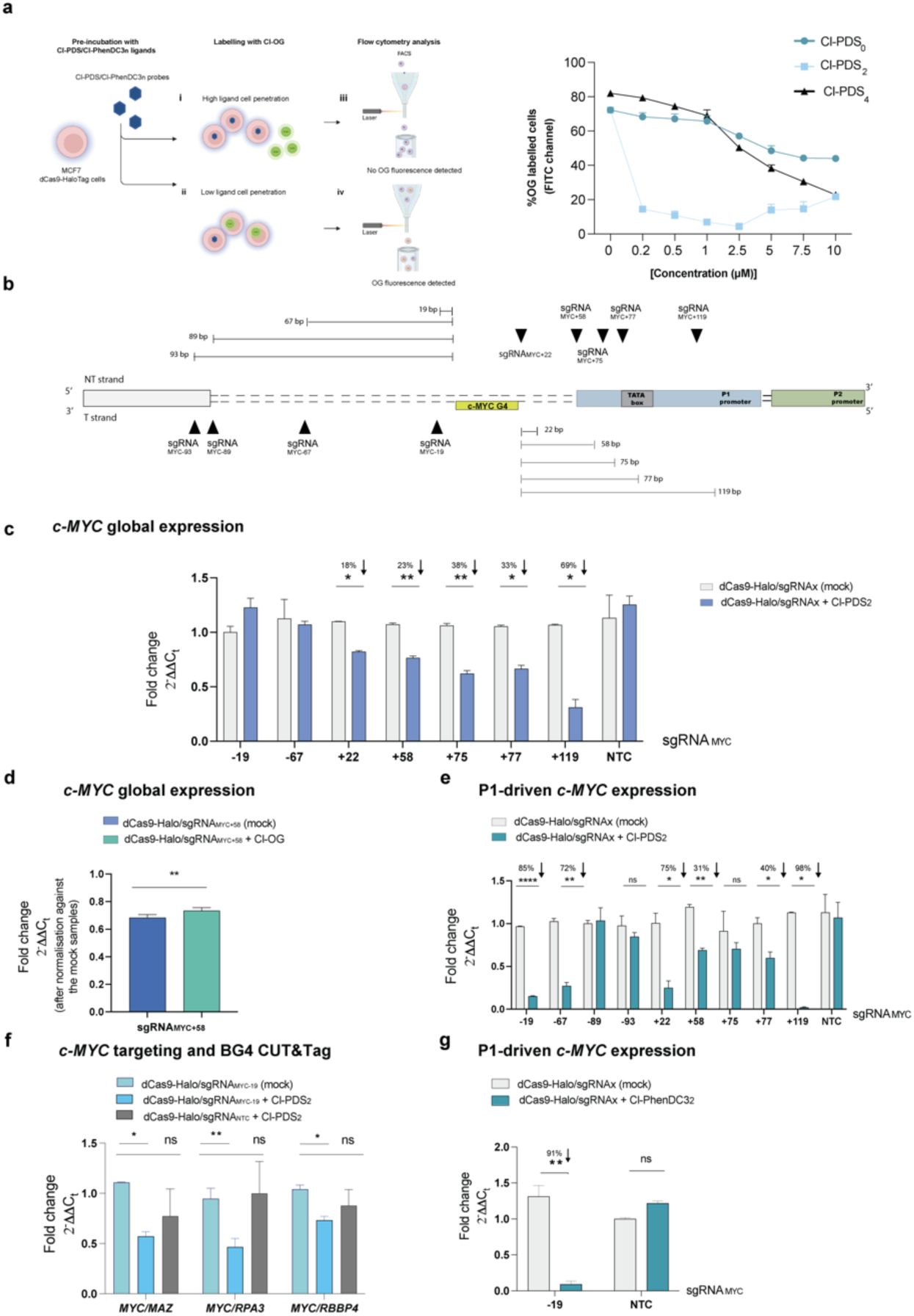
ATENA enables selective targeting of a G4 in the *c-MYC* promoter. **a**, (left) Schematic representation of the CAPA assay; (**i**) high probe cell penetration/reactivity results in Cl-OG inability to bind the HaloTag, conversely (**ii**) low probe cell penetration/reactivity allows Cl-OG to bind the HaloTag, resulting respectively in (**iii**) no Oregon Green fluorescence detected *via* Flow cytometry or (**iv**) presence of Oregon Green fluorescence signal in the FITC channel. (right) CAPA assay curves of Oregon Green fluorescence signal obtained upon treatment of MCF7 cells stably expressing dCas9-Halo with Cl-PDS_n_ followed by fluorophore incubation (n=2). Flow cytometry was performed using Attune NxT Flow cytometer (FITC channel), and data were then analyzed with FlowJo software. **b**, Schematic illustration of the *c-MYC* promoter with the annotated G4 (MYC-G4), sgRNA targeting region (black triangles), and their relative distance in bp from the MYC-G4. **c**, RT-qPCR for *c-MYC* expression in MCF7 cells stably expressing dCas9-Halo transfected with the indicated sgRNAs and incubated for 48h in the presence of (2.5 µM) Cl-PDS_2_ or DMSO (mock). The expression values are represented as fold change (2^-ΔΔCt^) with respect to the mock (DMSO-treated) and normalized for the housekeeping gene *GAPDH*. n=3, biological replicates, each with three technical replicates. **d,** RT-qPCR for *c-MYC* expression in MCF7 cells stably expressing dCas9-Halo transfected with sgRNA_MYC+58_ and incubated for 24h in the presence of either Cl-PDS_2_ (2.5 µM) or Cl-OG (5 µM). The expression values are represented as fold change (2^-ΔΔCt^) with respect to the mock (DMSO-treated) and normalized for the housekeeping gene *GAPDH*. n=2, biological replicates, each with two technical replicates. **e,** RT-qPCR for P1-driven *c-MYC* expression in MCF7 cells stably expressing dCas9-Halo transfected with the indicated sgRNAs and incubated for 48h in the presence of (2.5 µM) Cl-PDS_2_. The expression values are represented as fold change (2^-ΔΔCt^) with respect to the mock (DMSO-treated) and normalized for the housekeeping gene *GAPDH*. n=3, biological replicates, each with three technical replicates. **f,** BG4 CUT&Tag-qPCR for MCF7 cells stably expressing dCas9-Halo transfected with either sgRNA_MYC-19_ or sgRNA NTC and treated with DMSO (mock) or (2.5 µM) Cl-PDS_2._ BG4 accessibility was analyzed for *c-MYC* and normalized to three G4s in control gene sites (*RPA3*, *MAZ*, *RBBP4*). n=2, biological replicates, each with three technical replicates for BG4 and one for the negative (no BG4 treatment). **g,** RT-qPCR for P1-dependent *c-MYC* expression in MCF7 cells stably expressing dCas9-Halo transfected with sgRNA_MYC-19_ or sgRNA NTC and incubated for 48h in the presence of (2.5 µM) Cl-PhenDC3_2._ The expression values are represented as fold change (2^-ΔΔCt^) with respect to the mock (DMSO-treated) and normalized for the housekeeping gene *GAPDH*. n=3, biological replicates, each with two technical replicates. Data presented are the mean of n = number of independent biological samples. Statistical significance was calculated using a Welch-corrected two-tailed t-test in GraphPad Prism; p-value: ns > 0.05, * ≤0.05, ** ≤0.01, *** ≤0.001, **** ≤0.0001.

### Selective targeting of MYC-G4 through ATENA reveals no changes in *c-MYC* expression

After having identified conditions to decorate dCas9-Halo with G4-ligands in cells, we next set out to investigate transcriptional responses associated with individual G4-targeting using ATENA. We started our study by considering the G4 present in the promoter of the *c-MYC* proto-oncogene (MYC-G4), as this is one of the most studied and described G4s in the literature. Many studies have linked the targeting of MYC-G4 with ligands to transcriptional suppression of *c-MYC* ^12^. To assess the extent of transcriptional perturbation mediated exclusively by G4 stabilization, we designed a panel of sgRNAs to direct ATENA at MYC-G4.

Specifically, we designed sgRNAs targeting either the non-template strand (NT) bearing the G4-structure at its 3’ end or the opposite strand at its 5’ end (T), Fig. 2b. Based on our biophysical investigation, we reasoned that placing sgRNAs close enough to the G4 would have ensured G4-stabilization by G4-ligands tethered to dCas9-Halo (Fig. 1h). Considering PAM sequences available for dCas9-Halo binding at either G4 ends, we designed sgRNA_MYC-19_ and sgRNA_MYC-67_ that would place the protein complex on the T strand, respectively 19 and 67 base pairs away from the MYC-G4. Similarly, we generated sgRNA_MYC+22_ and sgRNA_MYC+58_, targeting the MYC-G4 from its 5’ end at a distance of 22 and 58 base pairs on the NT strand, respectively. To further investigate the optimal distance to achieve G4 stabilization in a cellular context, which might differ from our simple biophysical model, we have also designed sgRNA_MYC-89_ and sgRNA_MYC-93_, along with sgRNA_MYC+75_, sgRNA_MYC+77,_ and sgRNA_MYC+119_, also targeting the MYC-G4 at its 3’ and 5’ end, respectively, but at a further distance from the targeted structure (Fig. 2b). After cloning sequences encoding the various sgRNAs into a vector for mammal expression (see methods), we have transfected MCF7 cells stably expressing dCas9-Halo with individual sgRNAs, followed by treatment with either Cl-PDS_2_ or mock (DMSO) for 48 hours. We then measured changes in *c-MYC* expression using RT-qPCR, normalizing the expression level against individual samples transfected with the respective sgRNAs and mock-exposed. This enabled us to consider any change in gene expression potentially triggered by the positioning of the dCas9-Halo complex on the targeted site and, therefore, control for any transcriptional perturbation imposed by dCas9 not functionalized with ligands.

Unexpectedly, we observed that sgRNAs placing ATENA within a ∼50 bp window of the MYC-G4 on the T strand (sgRNA_MYC-19_, sgRNA_MYC-67_) in conjunction with Cl-PDS_2_ treatment led to negligible effects on the global expression of *c-MYC* (Fig. 2c), contrasting our biophysical predictions (Fig. 1h). Conversely when placing the complex on the NT strand with sgRNA_MYC+22_ and sgRNA_MYC+58_, we observed a modest decrease of *c-MYC* expression to 0.8-fold the mock (∼20% reduction), which could be indicative of G4-engagement of PDS mediated by ATENA (Fig. 2c). To investigate this further, we analyzed the transcriptional effects elicited by placing the ATENA further away from the MYC-G4 using sgRNA_MYC+75_, sgRNA_MYC+77,_ and sgRNA_MYC+119_, which should lead to abrogation of G4-engagement in a distance-dependent fashion, as observed in our biophysical measurements. To our surprise, we observed the opposite with *c-MYC* expression being drastically reduced the further away the complex was from the G4 (Fig. 2c), which cannot be consistent with a G4-mediated effect. By closer inspection of the promoter annotation, we noticed that sgRNA_MYC+58_ to sgRNA_MYC+119_ overlapped with the P1 promoter and TATA-box sequences. Since sgRNA_MYC+75_ and sgRNA_MYC+77_ are complementary to the region identifying a TATA box, we reasoned that ATENA occupies the TATA-box region and hinders transcription initiation, resulting in the observed *c-MYC* suppression. This effect is exacerbated when using sgRNA_MYC+119_ that targets the area next to the Transcriptional Starting Site (TSS) of the P1 promoter (4 bp downstream), further indicating a G4-independent transcriptional suppression.

Collectively, our results indicated that the reduction in *c-MYC* transcript levels is driven by ATENA’s interference with the TATA-box region, rather than by ligand-induced stabilization of the MYC-G4, suggesting that previous observations obtained with the equivalent of our sgRNA_MYC+58_ are likely affected by this^34^. To test this hypothesis further, we replaced the G4-stabilizer (Cl-PDS_2_) with a fluorophore (Cl-OG) and monitored *c-MYC* expression while using the sgRNA previously reported to provide the strongest downregulation (sgRNA_MYC+58_) ^34^. Under these conditions, we observed similar transcriptional downregulation of *c-MYC* compared to what was measured when treating with Cl-PDS_2_ (Fig. 2d), demonstrating that anchoring the dCas9 complex in the proximity of key transcriptional regions of the *c-MYC* promoter prevents any reliable evaluation of G4-mediated transcriptional effects. The interference of ATENA with *c-MYC* expression when placed in proximity to key promoter sites is consistent with what has been reported for CRISPRi studies^45,46^ and needs to be carefully considered when using dCas9-based strategies to target G4s^34^.

### ATENA confirms P1-dependent transcriptional expression associated with MYC-G4

It has been shown that multiple promoters globally contribute to regulating *c-MYC* expression ^47,48^. Therefore, we decided to examine how G4-targeting affects *c-MYC* expression regulated by specific promoters by analyzing transcripts originating from the two main ones: P1 and P2.

Indeed, it has been recently shown that genetic deletion of the MYC-G4 is associated with selective suppression of transcription from the P1 promoter and only a modest reduction of overall *c-MYC* expression, which is instead represented by the combined transcriptional output of both the P1 and P2 promoters ^20^. When using ATENA with Cl-PDS_2_ and monitoring P1-driven *c-MYC* expression, we observed a strikingly distance-dependent suppression of P1-mediated expression with sgRNA_MYC-19_ and sgRNA_MYC-67_. In particular, when using sgRNA_MYC-19_ we observed an 85% reduction of P1-mediated *c-MYC* expression (0.15-fold), whereas use of sgRNA_MYC-67_ led to a 72% reduction of the *c-MYC* expression (0.28-fold), Fig. 2e. Importantly, no significant changes in expression were detected when using ATENA in conjunction with sgRNA_MYC-89_ and sgRNA_MYC-93_ that place the Cl-PDS_2_ excessively distant from the targeted G4, which agrees with a G4-dependent transcriptional suppression (Fig. 2e). We then asked why the reduction in P1-driven expression observed with ATENA does not result in an overall decrease in *c-MYC* transcription. To achieve this, we also measured changes in transcription originating from the P2 promoter using promoter-specific qPCR primers (see methods). Notably, we observed an increase in P2-selective expression that suggests a compensatory mechanism activated by the cells in response to the G4-induced reduction of P1 transcription (Extended Data Fig. 3a), justifying the absence of statistically significant changes in global *c-MYC* expression detected when using primers amplifying regions common to both P1- and P2-derived transcripts.

Targeting the G4 from its 5’ end with sgRNA_MYC+22_ led to a 75% reduction of P1-driven *c-MYC* expression (0.25-fold) Fig. 2e. We also detected strong P1-dependent transcriptional repression when using sgRNA_MYC+58_ and sgRNA_MYC+75_, which might be affected by G4-independent transcriptional perturbation that we already observed when using these sgRNAs (Fig. 2e). Indeed, P1-mediated *c-MYC* expression was entirely abrogated when the ATENA was directed at sites overlapping close to the P1-TSS with sgRNA_MYC+119_ (Fig. 2e), which is consistent with G4-independent transcriptional inhibition. These observations further confirmed that using ATENA on the 5’ end of the MYC-G4 cannot reliably detect changes in gene expression that G4-targeting strictly mediates, and careful consideration of the promoter regulatory elements is needed.

Next, we sought to confirm that the observed changes in P1-driven *c-MYC* expression following treatment with sgRNA_MYC-19_ and sgRNA_MYC-67_ result specifically from ATENA-mediated targeting of the MYC-G4, rather than non-specific ligand interactions with other G4 structures. To address the potential for ligand-mediated off-target effects, we monitored the expression of *KRAS*, a gene known to contain a stable G4 structure in its promoter region. As shown in Extended Data Fig. 3b, directing ATENA specifically to the MYC-G4 did not alter *KRAS* expression, supporting the selectivity of ATENA-mediated G4 targeting. In contrast, free PyPDS treatment significantly lowered *KRAS* expression (0.20-fold), leading to an 86% transcriptional suppression (Extended Data Fig. 3c), which validates the ability of ATENA to confer G4-ligand selectivity towards individual G4s, whilst minimizing off-target effects.

### P1-dependent transcriptional inhibition is linked with protein displacement from MYC-G4

We further assessed the ability of ATENA to mediate selective MYC-G4 targeting by investigating perturbation in protein binding at MYC-G4 upon ligand stabilization. It has been proposed that ligands bound to G4s can displace key transcription factors and regulatory proteins, leading to the observed transcription suppression ^21,49^. Therefore, we reasoned that if ATENA was correctly positioned to enable ligand-G4 interaction, we should have observed reduced protein accessibility to the G4. To measure this, we used the G4-selective antibody BG4 ^4^ and performed CUT&Tag ^7^ coupled with qPCR to compare the efficiency of BG4 in enriching for MYC-G4, targeted by ATENA, against 3 independent validated G4-sites (MAZ, RPA3, RBBP4) that should not be affected as they are not targeted by ATENA. This enabled us to assess relative protein accessibility at these individual G4 sites under different conditions, as previously described ^20^. As displayed in Fig. 2f, when treating cells with Cl-PDS_2_ using sgRNA_MYC-19_, we observed a consistent reduction of BG4 signal that was not detected for the Non-Targeting Control (NTC), irrespective of the reference G4 used. This result strongly suggests that ATENA can be used to guide Cl-PDS_2_ selectively to the MYC-G4 structure, resulting in a decrease in binding of the BG4 antibody to MYC-G4 due to the binding competition of the ligand that leads to displacement of the antibody from the G4. This supports a model in which ligand-mediated G4 stabilization suppresses P1-driven *c-MYC* transcription by outcompeting binding of transcriptional effectors at the MYC-G4^21^.

### P1-dependent transcriptional suppression is recapitulated with PhenDC_3_

To further validate our findings, we used ATENA to deploy a different G4 ligand - PhenDC3 ^31^ - to stabilize the MYC-G4. The PEG2-functionalized analogue, Cl-PhenDC3_2_, also efficiently labeled dCas9–Halo in cells, as confirmed by CAPA (Extended Data Fig. 2d). When using ATENA with Cl-PhenDC3_2_ and sgRNA_MYC-19_, we observed a significant inhibition of P1-mediated *c-MYC* transcription (0.09-fold), leading to a 91% reduction of expression (Fig. 2g), which is even greater than what observed with Cl-PDS_2_, likely reflecting the stronger G4 stabilization capacity of Cl-PhenDC3_2_, as indicated by biophysical CD measurements (Extended Data Table 4). This result suggests that ATENA can also be leveraged to compare the relative potency and biological impact of different G4 ligands when deployed at the same genomic target.

Collectively, our data indicate that ATENA can successfully target MYC-G4 in a ligand-independent fashion, leading to a detectable G4-engagement and corresponding P1-specific *c-MYC* suppression, validating recent findings generated by genetic deletion of the sequence responsible for the MYC-G4 folding^20^.

### The MYC-G4 selective molecule DC-34 validates ATENA

Encouraged by the P1-driven *c-MYC* suppression observed upon targeting MYC-G4 with ATENA, we explored whether a similar phenotype could be elicited when using ligands that display some inter-G4 selectivity. To this end, we leveraged the MYC-G4 selective ligand DC-34 (Fig. 3a), which shows a strong binding affinity for the MYC-G4 with high selectivity over other G4 structures such as *KRAS* and *c-KIT* ^24^. We treated MCF7 cells with increasing concentrations of DC-34 for 48 hours before measuring changes in *c-MYC* expression by RT-qPCR. As observed with ATENA, treatment with DC-34 caused negligible dose-dependent changes in global *c-MYC* expression (Fig. 3b), supporting the notion that selective targeting of MYC-G4 does not impact the overall expression of *c-MYC* in MCF7 cells. Conversely, when measuring P1-mediated transcription, DC-34 revealed a strong dose-dependent suppression that plateaued at 74% reduction when treating with 10 μM of the ligand, (0.26-fold), Fig. 3c. This indicated that using an inter-G4 selective ligand for selective MYC-G4 targeting led to observations comparable to those obtained using ATENA, further corroborating the validity of our platform for single G4-targeting.

**Figure 3:**
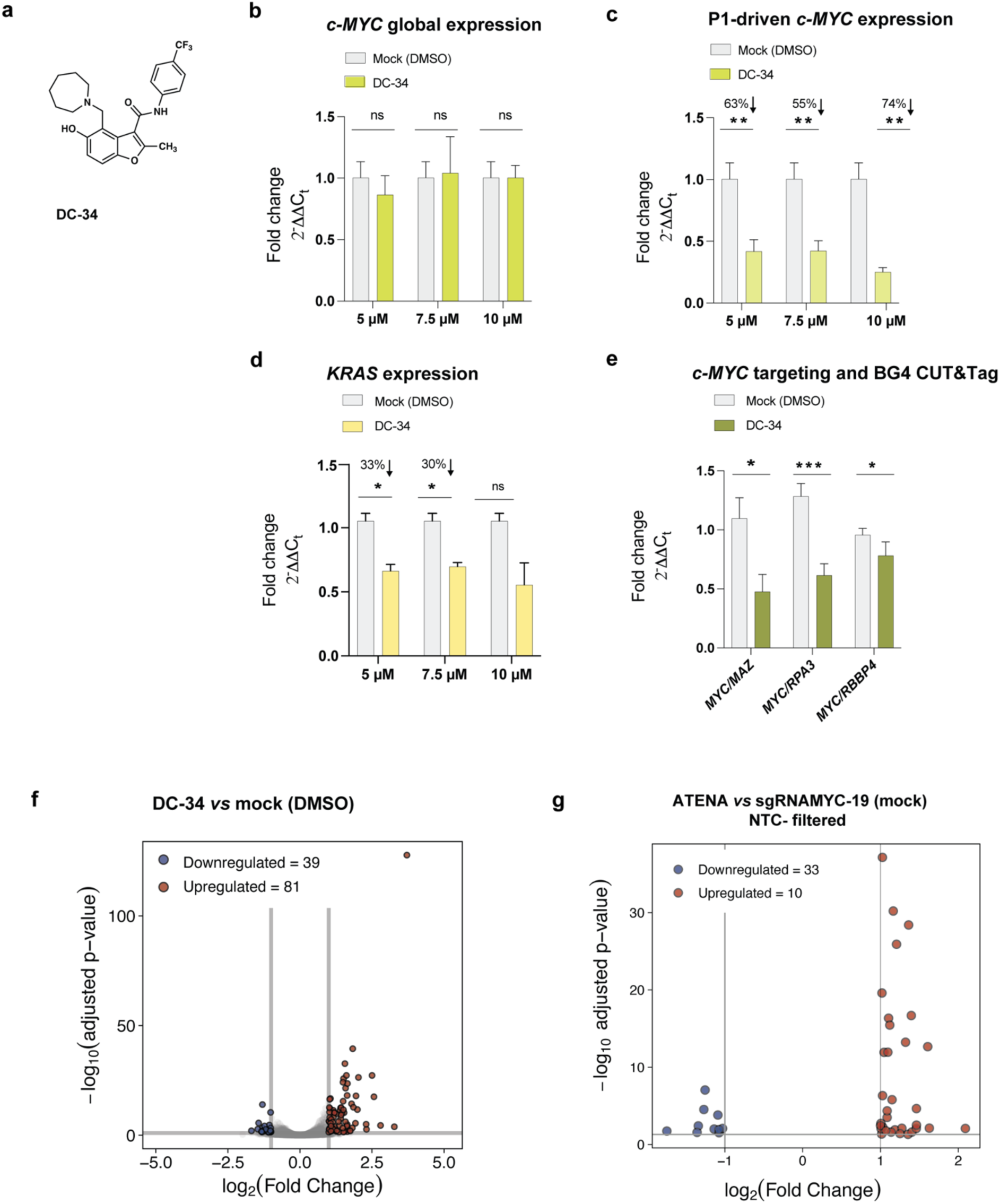
The selective G4-MYC ligand DC-34 matches ATENA. **a**, Chemical structure of DC-34. **b,** RT-qPCR for *c-MYC* expression in MCF7 cells treated with increasing concentration of DC-34 for 48h. Values are represented as fold change (2^-ΔΔCt^) with respect to the mock (DMSO-treated) and normalized for the housekeeping gene *GAPDH*. n=3, biological replicates, each of which included two technical replicates. **c**, RT-qPCR for P1-driven *c-MYC* expression in MCF7 cells treated with different concentrations of DC-34 for 48h. Values are represented as fold change (2^-ΔΔCt^) with respect to the mock (DMSO-treated) and normalized for the housekeeping gene *GAPDH*. n=3, biological replicates, each of which included two technical replicates. **d**, Evaluation of potential DC-34 off-targets by analyzing the *KRAS* expression using the same samples reported in c. **e,** BG4 CUT&Tag-qPCR for MCF7 cells treated with DMSO (mock) or (7.5 µM) DC-34. BG4 accessibility was analyzed for *c-MYC* and normalized to three G4s in control gene sites (*RPA3*, *MAZ*, *RBBP4*). n=2, biological replicates, including three technical replicates for BG4 and one for the negative (no BG4 treatment). **f,** Volcano plot of DEGs in MCF7 cells treated with DC-34 *vs* cells treated with DMSO (mock). Gray = non-significant genes (FDR ≥ 0.05 or |log₂FC| < 1); red = up-regulated DEGs (FDR < 0.05, log₂FC ≥ +1); blue = down-regulated DEGs (FDR < 0.05, log₂FC ≤ −1).**g,** Volcano plot of DEGs in MCF7 cells transfected with sgRNA_MYC-19_ treated with Cl-PDS_2_ *vs* mock, after sgRNA NTC filtering; Red = up-regulated DEGs (FDR < 0.05, log₂FC ≥ +1); blue = down-regulated DEGs (FDR < 0.05, log₂FC ≤ −1). Plot includes only genes passing FDR < 0.05 and |log₂FC| ≥ 1. The data presented are the mean of n = number of independent biological samples. Statistical significance was calculated using a Welch-corrected two-tailed t-test in GraphPad Prism; p-value: ns > 0.05, * ≤0.05, ** ≤0.01, *** ≤0.001, **** ≤0.0001.

Next, we assessed the selectivity of DC-34 for MYC-G4 relative to another G4-containing promoter, as we have done for ATENA. Our observations indicated that ATENA did not lead to detectable changes in *KRAS* expression when guided to MYC-G4 (Extended Data Fig. 3b). In contrast, treatment with DC-34 resulted in a dose-dependent reduction in *KRAS* expression (∼30%, Fig. 3d), which was less pronounced than what was observed for *c-MYC* under the same conditions, indicating that MYC-G4 is the primary target of DC-34, but residual off-target binding to other G4s may occur at high concentrations. This likely reflects the structural similarity shared between different G4s, which substantially complicates inter-G4 selective targeting with small-molecule ligands.

To further confirm that DC-34 downregulates *c-MYC* through direct G4 binding, we also performed the BG4 CUT&Tag qPCR upon ligand treatment that we optimized for ATENA to assess protein occupancy upon treatment. As shown in Fig. 3e, DC-34 treatment significantly reduced the BG4 signal at MYC-G4, which is consistent with the decreased protein accessibility induced by MYC-G4 selective targeting with ATENA. This further supports the use of BG4 CUT&Tag qPCR as an indirect measure of ligand-mediated G4 stabilization associated with transcriptional suppression.

### Transcriptome-wide comparison of ATENA with DC-34

To further assess the inter-G4 selectivity provided by ATENA and DC-34, we analyzed transcriptome-wide gene-expression changes through mRNA-seq. Specifically, we generated mRNA-Seq datasets for MCF7 cells treated with either DC-34 or ATENA (Cl-PDS_2_ in conjunction with sgRNA_MYC-19_) and compared those to transcriptome-wide changes induced by the generic G4-ligand PyPDS. We hypothesized that the inter-G4 selectivity provided to Cl-PDS_2_ by ATENA should be reflected by a substantially lower number of differentially expressed genes (DEGs) when compared to free PyPDS. Indeed, treatment of MCF7 cells with PyPDS (2.5 μM) for 6 hours altered the expression of 2,228 genes (319 downregulated, 1907 upregulated, FDR<0.05, Extended Data Fig. 4). In contrast, treatment with the MYC-G4 selective ligand DC-34 (7.5 μM, 6h) affected only the expression of 120 genes (39 downregulated, 81 upregulated; FDR<0.05, Fig. 3f), indicating that the enhanced inter-G4 selectivity of DC-34 is translated into a lower number of genes being differentially expressed when compared to PyPDS. Additionally, mRNA-Seq confirmed that DC-34 did not significantly alter *c-MYC* expression globally, as already observed in our RT-qPCR data, further indicating the selective downregulation of P1 transcripts when targeting MYC-G4, an essential consideration when devising G4-based therapies aiming at suppressing *c-MYC* expression. Given the substantial reduction in DEGs observed with DC-34 relative to a global G4-stabilizer like PyPDS, we hypothesized that conjugation of Cl-PDS_2_ to dCas9 in ATENA should also reduce the extent of transcriptional perturbations. To investigate this, we have transfected MCF7 cells with sgRNA_MYC-19_ and treated them with Cl-PDS_2_ (6 h) before performing mRNA-Seq.

Initially, we performed a standard differential expression analysis comparing ATENA (sgRNA_-MYC-19_ or sgRNA NTC + Cl-PDS_2_) to their DMSO-treated controls using DESeq2. We filtered the resulting DEGs for statistical significance (adjusted p-value < 0.05) and magnitude of change (|log2FoldChange| ≥ 1), Table S9.

However, to identify genes uniquely responsive to MYC-targeted ligand recruitment, we then excluded all genes that were also differentially expressed in the sgRNA NTC + Cl-PDS_2_ *vs* DMSO comparison, using the same filtering criteria. This filtering allowed us to isolate transcriptional effects that were specific to the combination of the sgRNA_MYC-19_ and the ligand, rather than shared responses with sgRNA NTC that reflect more unspecific effects of the dCas9 platform (Fig. 3g). This analysis identified only 43 DEGs, underscoring the exceptional specificity conferred to Cl-PDS_2_ when conjugated to dCas9 as opposed to the free PyPDS ligand.

To evaluate if ATENA, DC-34, and free PyPDS shared any of the transcriptional changes elicited, we compared DEGs observed in: DC-34 vs ATENA (NTC-filtered), PyPDS vs ATENA (NTC-filtered), and PyPDS vs DC-34, while also testing for enrichment in *c-MYC*-related pathways^50^. When comparing DC-34 *vs* ATENA (NTC-filtered), and free PyPDS *vs* ATENA (NTC-filtered), we found an overlap of a few genes that were not enriched in any pathway, suggesting that these approaches yield largely orthogonal transcriptomic profiles, consistent with their distinct mechanisms of delivery and engagement (Tables S10-11). Finally, when comparing PyPDS vs DC-34, we observed 24 shared DEGs (Table S12). However, these DEGs failed to enrich for any known pathways or MYC-related functions, indicating that even structurally distinct G4 ligands with overlapping target preferences can elicit unique transcriptional responses, likely due to differences in binding affinity, cellular uptake, and selectivity.

Altogether, these analyses indicated that ATENA, DC-34, and PyPDS can elicit distinct transcriptional responses, with negligible overlap and no shared enrichment for *c-MYC*-related genes. This reinforces the conclusion that ATENA - by spatially confining ligand activity to a single G4 at the *c-MYC* promoter - induces highly selective gene expression changes, contrasting with the broader, less discriminating effects of freely diffusing G4-ligands. Additionally, our mRNA-Seq analysis further confirmed that MYC-G4 targeting is not associated with significant *c-MYC* downregulation in MCF7 cells, and caution is needed when designing MYC-based therapeutics based on G4-targeting.

### Targeting of the *c-MYC* i-motif with ATENA is associated with transcriptional stimulation

After validating the suitability of ATENA for the selective targeting of individual G4-structures in the genome - exemplified by the MYC-G4 - we next explored whether this platform could be adapted to interrogate other DNA secondary structures, such as i-motifs (iMs). Similarly to G4s, iMs are stabilized by Hoogsteen hydrogen bonding. However, they form in cytosine-rich regions of the genome, typically complementary to G-rich G4-forming sequences^51^. The formation of iMs in cells has been recently validated in living cells by both immunofluorescence^23^ and genome-wide mapping^22^. Like G4s, iMs have been implicated in transcriptional regulation^35^, although their mechanistic roles remain less well characterized compared to G4s.

To assess the potential of individual iMs to modulate gene expression, we decided to use ATENA in combination with selective iM ligands. Conveniently, the *c-MYC* promoter also bears an iM structure complementary to the G-rich sequence forming the MYC-G4^52^, which has been reported to modulate *c-MYC* expression when targeted with selective iM ligands^53–55^. We therefore reasoned that the same sgRNAs optimized for selective targeting of MYC-G4 could be repurposed to localize ATENA to the MYC-iM, enabling selective iM targeting when decorated with an appropriate ligand. To this end, we utilized a recently developed class of short peptides from the Waller group, which show high affinity and specificity for iMs over G4s^33^. Among those, we selected the RVS peptide (pep-RVS) for synthetic ease and potential amenability for modification. To enable compatibility with ATENA, we chemically modified pep-RVS with chloroalkene side chains to ensure covalent attachment onto the Halo-Tag (SI-2 and SI-7), generating Cl-pep-RVS_n_ analogs with different PEG-linkers (n). We first confirmed that functionalization with chloroalkene did not affect the ability of pep-RVS to bind iMs *via* UV titrations, confirming that it bound the c-MYC i-motif structure with a *K*_d_ of 0.35 ± 0.12 µM compared to >33 µM for G4 (Fig. SI-18). We next assessed the ability of Cl-pep-RVS_n_ analogs to covalently bind to the dCas9-Halo in living cells using CAPA. As shown in Extended Data Fig. 5a, all Cl-pep-RVS_n_ analogs displayed good cellular permeability and effectively labelled dCas9-Halo at low µM concentrations. However, Cl-pep-RVS_4_ performed best in CAPA, providing a CP_50_ value of 2.3 µM, and was selected for further application in ATENA.

To evaluate whether targeting the MYC-iM affects *c-MYC* transcription, we employed ATENA with sgRNA_MYC-19_ and treated the cells with Cl-pep-RVS_4_ for 48h, using the same conditions previously optimized for G4 targeting. Notably, ATENA-mediated iM-targeting led to a significant increase of P1-driven (∼2-fold increase) *c-MYC* transcription (Fig. 4b), in contrast to the transcriptional repression observed upon MYC-G4 targeting. Interestingly, this P1-mediated upregulation was accompanied by a decrease in P2-driven transcription (Fig. 4b), resulting in minimal net changes in global *c-MYC* expression - mirroring the outcome observed when targeting the G4 motif.

**Figure 4:**
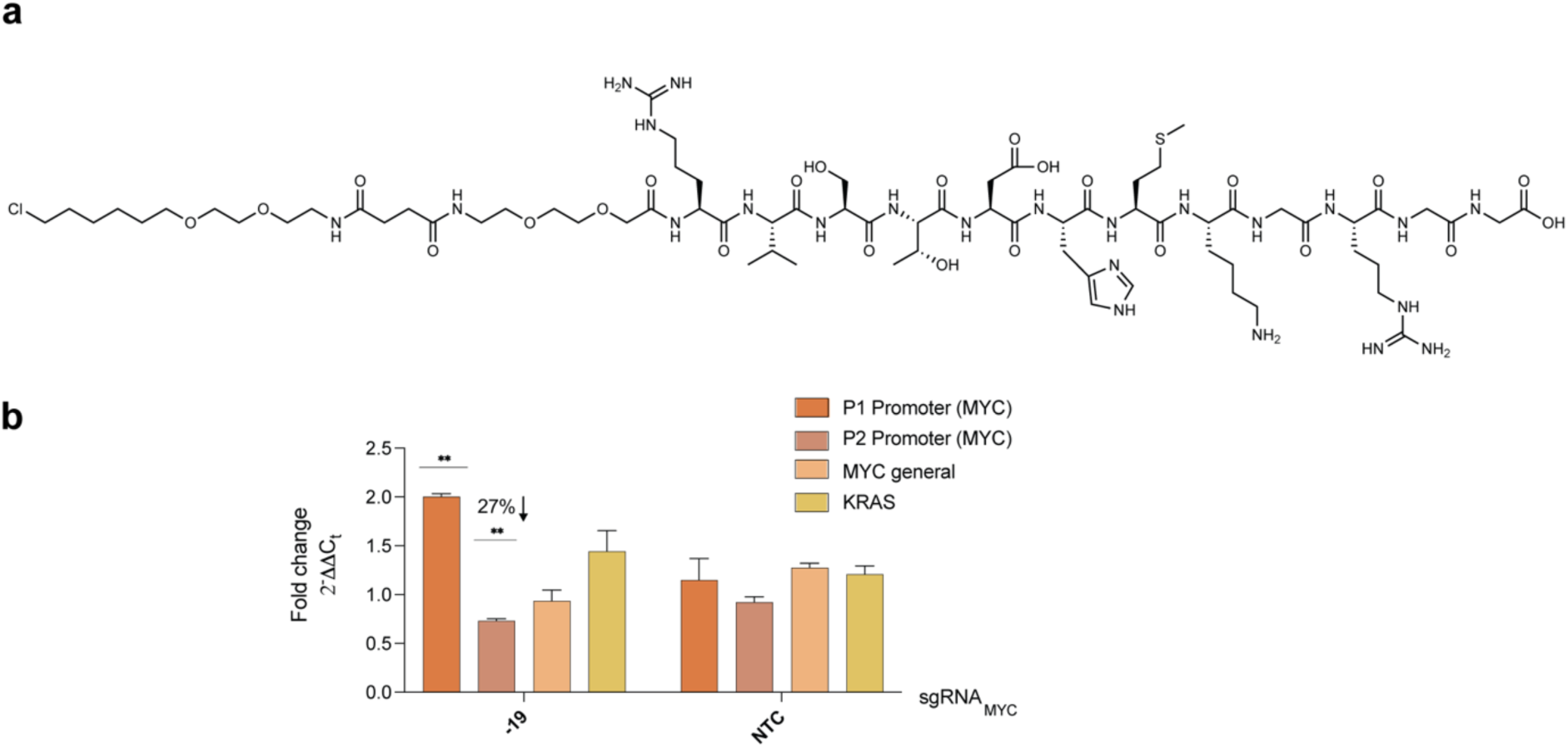
ATENA and *c-MYC* iM targeting. **a,** Chemical structure of chloroalkane-modified pep-RVS with a 2-PEG linker (Cl-pep-RVS_2_). **b,** RT-qPCR of the indicated genes in MCF7 cells stably expressing dCas9-Halo transfected with sgRNA_MYC-19_ or sgRNA NTC and incubated for 48h in the presence of 10 µM of Cl-pep-RVS2 or DMSO (mock). The expression values are represented as fold change (2^-ΔΔCt^) with respect to the mock (DMSO-treated) transfected samples and after normalization for the housekeeping gene (GAPDH). n=2, biological replicates, each with three technical replicates. The data presented are the mean of n = number of independent biological samples. Statistical significance was calculated using a Welch-corrected two-tailed t-test in GraphPad Prism; p-value: ns > 0.05, * ≤0.05, ** ≤0.01, *** ≤0.001, **** ≤0.0001.

Overall, these results underscore the modularity of ATENA in targeting distinct DNA secondary structures within the same genomic locus simply by varying the conjugated ligand. By exploiting the high iM selectivity of pep-RVS, we show that selective iM targeting is associated with P1-dependent transcriptional activation of *c-MYC -* a functional effect opposite to that seen with G4 engagement. This highlights the capacity of ATENA to disentangle the complex regulatory roles of overlapping secondary structures in gene promoters.

### PDS can act as a molecular glue of specific G4-protein interactions

After establishing the reliability of ATENA to accurately measure biological responses mediated by individual DNA secondary structures, using *c-MYC* as a case study, we next questioned if this platform could be expanded to study transcriptional responses that are uniquely associated with specific ligands. For instance, previous reports indicated that PDS treatment of MCF7 cells led to significant upregulation, rather than repression, of the long non-coding RNA *PVT1*-an observation we confirmed also for PyPDS by RT-qPCR and mRNA-seq (Extended data Fig. 3a) ^43^. Therefore, we sought to leverage ATENA to determine if PDS-mediated *PVT1* upregulation was a direct response to the specific stabilization of the G4 in its promoter or rather an indirect effect caused by global G4 stabilization. Previous CUT&Tag experiments performed in MCF7 cells identified a clear G4-peak in the *PVT1* promoter, ^56^ which we have used to design sgRNAs for ATENA-based targeting of the PVT1-G4. Considering available PAM sequences, we designed two sgRNAs targeting the PVT1-G4 at either 20 base pairs from its 5’ end (sgRNA_PVT1-20_) or 33 base pairs from its 3’ end (sgRNA_PVT1+33_) and that display no overlap with any annotated regulatory site (Fig. 5a). Upon transfection with either sgRNA_PVT1-20_ or sgRNA_PVT1+33_ and treatment with Cl-PDS_2_, we observed a boosted *PVT1* expression of ∼4-fold, consistent with observations reported using free PDS (Fig. 5b). G4-ligands typically compete with regulatory proteins for G4-binding, which leads to transcriptional suppression, as we recapitulated measuring BG4 occupancy upon MYC-G4 targeting (Fig. 2f). Therefore, we questioned whether PDS could instead act as a molecular glue when binding to the PVT1-G4, leading to enhanced protein binding to the PVT1-G4 and, thus, justifying the observed transcriptional increase upon PDS treatment. We performed BG4 CUT&Tag qPCR on PVT1-G4 target by ATENA and in conjunction with Cl-PDS_2_ and observed a rise in BG4 occupancy at *PVT1* of ∼2-fold (Fig. 5c). This contrasts the transcriptional response elicited by the same ligand when targeting a different G4 (MYC-G4, Fig. 2f) and points to the role of PyPDS as a molecular glue for protein-G4 interactions within the PVT1-G4. Moreover, this observation suggests that the previously described increase in *PVT1* expression elicited by PyPDS treatment in MCF7 cells reflects the genuine response of the ligand targeting the PVT1-G4 and cannot be ascribed to a secondary response associated with global G4-targeting.

**Figure 5:**
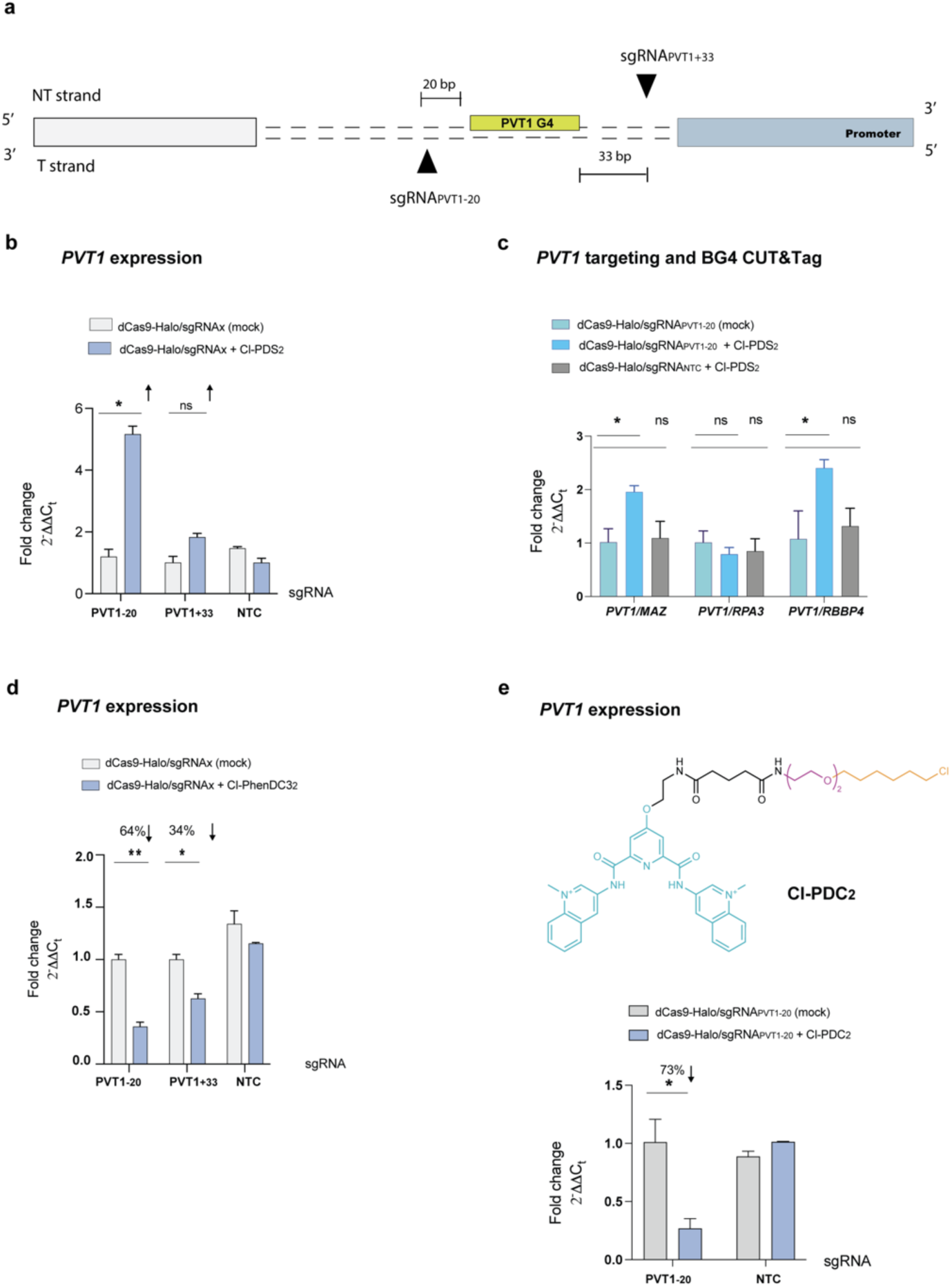
ATENA unveils a ligand-dependent transcriptional response of the *lncPVT1*. **a**, Schematic overview of the *PVT1* promoter containing annotation of the predicted G4, sgRNA targeting region (black triangles) and their relative distance in bp from the G4 forming sequence **b,** RT-qPCR for *PVT1* expression in MCF7 cells stably expressing dCas9-Halo, transfected with either sgRNA_PVT1-20_, sgRNA_PVT1+33_ or sgRNA NTC and treated with (2.5 µM) Cl-PDS_2_ for 48h after transfection. The expression values are represented as fold change (2^-ΔΔCt^) with respect to the mock (DMSO-treated) and normalized for the housekeeping gene *GAPDH*; n=3, biological replicates, each of which included two technical replicates. **c,** BG4 CUT&Tag-qPCR for MCF7 cells stably expressing dCas9-Halo transfected with either sgRNA_PVT1-20_ or sgRNA NTC and treated with DMSO (mock) or (2.5 µM) Cl-PDS_2._ BG4 accessibility was analyzed for *PVT1* and normalized to three G4s in control gene sites (*RPA3*, *MAZ*, *RBBP4*). n=2, biological replicates, including three technical replicates for BG4 and one for the negative (no BG4 treatment). **d**, RT-qPCR for *PVT1* expression in MCF7 cells stably expressing dCas9-Halo, transfected with either sgRNA_PVT1-20_, sgRNA_PVT1+33_ or sgRNA NTC and treated with (2.5 µM) Cl-PhenDC3_2_ for 48h after transfection. Values are represented as fold change (2^-ΔΔCt^) with respect to the mock (DMSO-treated) and normalized for the housekeeping gene *GAPDH*. n=3, biological replicates, each of which includes two technical replicates. **e,** (Top) Chemical structure of chloroalkane-modified PDC (Cl-PDC_2_) with the PEG linker length of two. **e**, (bottom) RT-qPCR for *PVT1* expression in MCF7 cells stably expressing dCas9-Halo, transfected with either sgRNA_PVT1-20_ or sgRNA NTC and treated with (2.5 µM) Cl-PDC_2_ for 48h after transfection. Values are represented as fold change (2^-ΔΔCt^) with respect to the mock (DMSO-treated) and normalized for the housekeeping gene *GAPDH*. n=3, biological replicates, each of which includes two technical replicates. Statistical significance was calculated using a Welch-corrected two-tailed t-test in GraphPad Prism; p-value: ns > 0.05, * ≤0.05, ** ≤0.01, *** ≤0.001, **** ≤0.0001.

We next asked whether the increase in *PVT1* expression measured with PyPDS was limited to this molecule or if a more general response could be observed with any G4-ligand. MCF7 treatment with free PhenDC3 led to the suppression rather than the enhancement of *PVT1* expression, suggesting that different G4-ligands might elicit different responses when targeting an identical G4 (Extended Data Fig. 5b).

To further investigate this, we targeted selectively the PVT1-G4 using ATENA in conjunction with Cl-PhenDC3_2_ and sgRNA_PVT1-20_ or sgRNA_PVT1+33_. Under these conditions, we observed a suppression of *PVT1* transcription, 64% and 34%, respectively, (expression level of the sample was 0.36-fold and 0.66-fold the mock respectively), (Fig. 5d), which is consistent with free PhenDC3 treatment and opposite to what was observed with PyPDS (Extended Data Fig. 5c). This suggested that G4-ligands can elicit a different response when bound to the same G4, possibly reflecting binding modalities that can either increase or prevent protein accessibility, which can be indirectly measured by BG4 occupancy (Extended Data Fig. 5d). This is an essential factor to consider when using different G4-ligands to infer the biology associated with these secondary structures, as it is often assumed that G4 ligands will all behave the same.

To further investigate these ligand-specific observations, we have also synthesized a chloroalkene analog of a third widely used G4-ligand: Pyrido Dicarboxamide (PDC, Fig. 5e) ^57^. We functionalized the PDC scaffold with a PEG2 chloroalkane side chain (Cl-PDC_2_) to mimic the PyPDS and PhenDC3 analogues used in ATENA (Fig. 4e, SI-1). After validating the labelling efficiency of Cl-PDC_2_ through CAPA (Fig. SI-5), we used it in conjunction with sgRNA_-20_ to target the *PVT1* promoter and investigate associated transcriptional responses. Similarly, to what was observed with PhenDC3, PDC lowered *PVT1* expression (0.27-fold), leading to a 73% reduction (Fig. 5e), suggesting that PDC interacts with the PVT1-G4 in a manner reminiscent of PhenDC3 and causes protein displacement from the G4. Structurally, the PDC scaffold is indeed similar to PhenDC3, displaying methylated nitrogens on the quinolines that are facing opposite orientation compared to PDS (Fig. 5e). Moreover, both PhenDC3 and PDC lack the amino-side chains present in PyPDS, further highlighting the structural similarity between these two scaffolds, which might recapitulate the similar response observed. Collectively, our results indicated that the transcriptional responses elicited by ligands at individual G4s depend heavily on the structural nature of the ligand and on their binding modality, which might lead to protein displacement at the G4-site or might act as a molecular glue enhancing G4-protein interactions. This strongly suggests that transcriptional changes observed upon G4-ligand treatment should be interpreted as ligand-specific outcomes, reflecting the response to a specific ligand at a specific G4 structure rather than the endogenous function of the DNA structure.

### Targeting cell-specific G4s with ATENA reveals transcription-dependent response to ligands

CUT&Tag and other chromatin-compatible G4-mapping methods, such as BG4-ChIP and Chem-Map, have shown that genomic G4 distribution is cell-specific and predominantly located at promoters of highly expressed genes ^7,9,10,58^. Therefore, we decided to investigate biological responses attained when directing a ligand towards previously unexplored MCF7-specific G4s. Specifically, we aimed to determine whether the biological relevance of individual G4s and their response to ligand binding correlate with the expression levels of the associated genes. We leveraged the existing dataset on G4-distribution in MCF7 cells previously obtained using CUT&Tag^56^. This dataset identified a G4-peak in the promoter of the highly expressed *HMGN1* gene - encoding for a non-histone chromosomal protein able to interact with nucleosomes and regulate chromatin structure ^59,60^ - as unique to MCF7 cells compared to other cell lines^56^. We designed sgRNA_HMGN1-22_ and sgRNA _HMGN1+34_ to target the HMGN1-G4 at 22 and 34 base pairs, respectively, at its 3’ and 5’ ends, ensuring no overlap with known functional regions (Fig. 6a). After transfecting MCF7 cells expressing dCas9-Halo with sgRNA _HMGN1-22_ and sgRNA _HMGN1+34_, we incubated them with Cl-PDS_2_ for 48 hours, as per the optimized ATENA protocol. Under these conditions, we measured a striking 99% reduction of *HMGN1* expression (0.01-fold) when targeting its G4 at the closest distance of 22 base pairs with sgRNA_HMGN-22_ (Fig. 5b). *HMGN1* downregulation was partially attenuated when placing ATENA further away from the G4 with sgRNA_HMGN+34_, consistent with the distance-dependent ligand engagement observed for other G4s (Fig. 6b). To place the observed *HMGN1* down-regulation in a biologically meaningful context, we examined the role of this gene in breast-cancer dormancy - an epigenetic-driven, non-replicative state, from which cancer cells “awaken”, causing cancer relapse and resistance to therapy. In dormant MCF7 cells (estrogen-deprived), we inspected the epigenetic changes, chromatin accessibility, and transcriptional profile at the *HMGN1* promoter. Dormant cells exhibited loss of the active histone mark H3K4me3, gain of the repressive histone mark H3K27me3, and a corresponding drop in *HMGN1* expression; these epigenetic and transcriptomic changes were partially reversed upon cell-cycle re-entry - “awakening”^61^ (Extended data Fig. 6a, 6b). Remarkably, ATENA-mediated stabilization of the HMGN1-G4 reproduced this repressive transcriptional state (Fig. 6b). Therefore, these convergent observations suggest that the HMGN1-G4 could act as an epigenetic switch: stabilizing the structure could indeed reinforce the *HMGN1* repressive state characteristic of dormancy and might be investigated as a strategy to maintain residual tumor cells in a dormant state.

**Figure 6:**
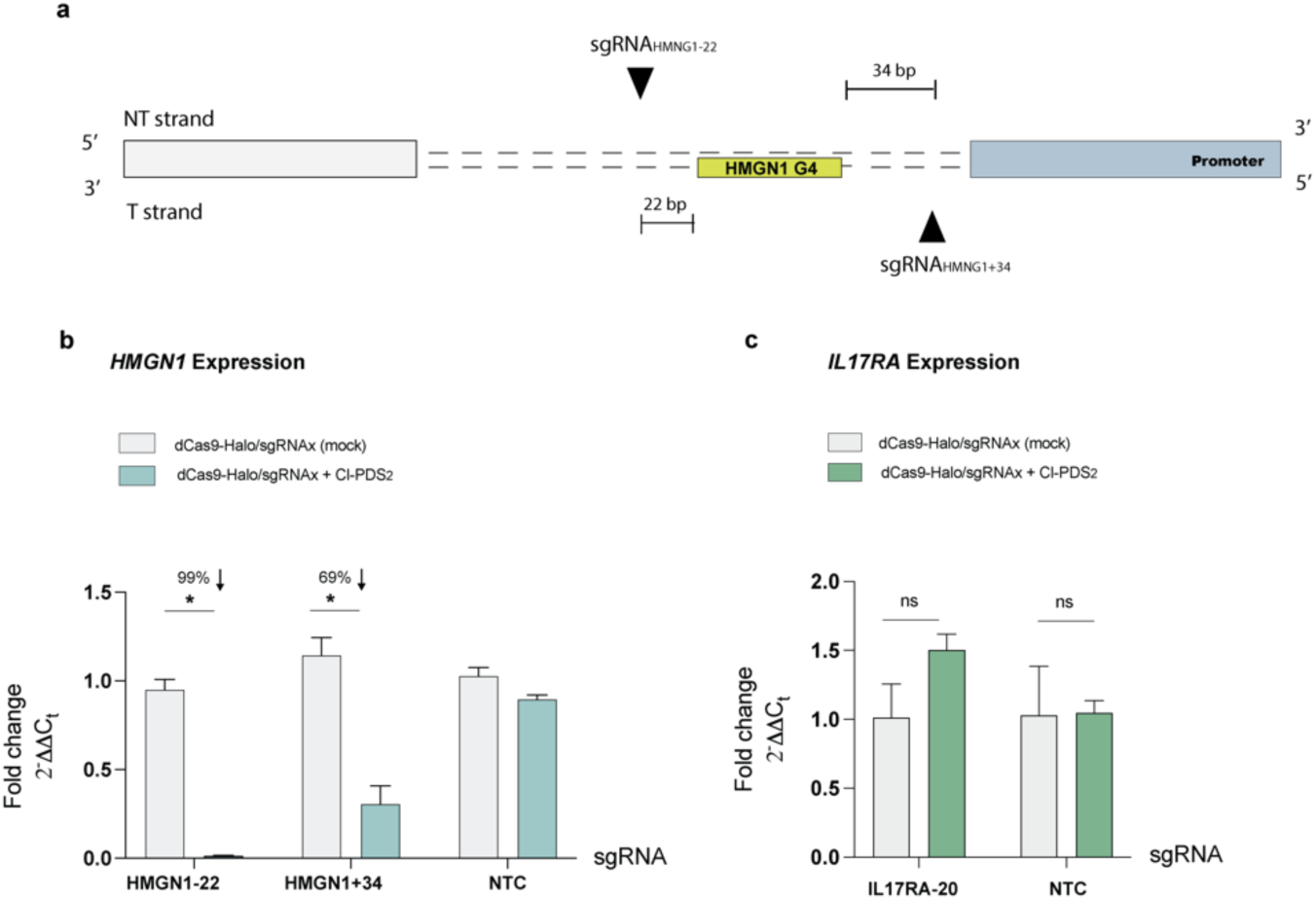
Targeting of *de novo* G4s with ATENA uncovers a transcriptionally dependent functional response. A schematic overview of the *HMGN1* promoter contains annotation of the predicted G4, two sgRNA designed to target HMGN1-G4 (black triangles), and their relative distance in bp from the G4-forming sequence. **b,** RT-qPCR for *HMGN1* expression in MCF7 cells stably expressing dCas9-Halo, transfected with either sgRNA_HMGN1-22_, sgRNA_HMGN1+34_ or sgRNA NTC and treated with (2.5 µM) Cl-PDS_2_ for 48h after transfection. The expression values are represented as fold change (2^-ΔΔCt^) with respect to the mock (DMSO-treated) and normalized for the housekeeping gene *GAPDH*; n=2, biological replicates, each of which includes two technical replicates. **c**, RT-qPCR for *IL17RA* expression in MCF7 cells stably expressing dCas9-Halo, transfected with either sgRNA_IL17RA-20_ or sgRNA NTC and treated with (2.5 µM) Cl-PDS_2_ for 48h after transfection. The expression values are represented as fold change (2^-ΔΔCt^) with respect to the mock (DMSO-treated) and normalized for the housekeeping gene *GAPDH*; n=2, biological replicates, each of which includes two technical replicates. Data presented are the mean of n = number of independent biological samples. Statistical significance was calculated using a Welch-corrected two-tailed t-test in GraphPad Prism; p-value: ns > 0.05, * ≤0.05, ** ≤0.01, *** ≤0.001, **** ≤0.0001.

Altogether, these findings indicated that targeting a cell-specific G4 located in the promoter of a highly transcribed gene can profoundly suppress gene expression, suggesting that maintaining G4-homeostasis at the promoter of highly transcribed genes is key to sustaining elevated expression levels and making these G4s particularly sensitive to ligands. Therefore, the varied transcriptional response observed upon G4-ligand treatment might reflect the relative relevance of different G4s in sustaining gene expression in the specific cell line studied.

To corroborate this hypothesis, we used ATENA to target a G4 present in the promoter region of a gene expressed at very low levels in MCF7 cells: *IL17RA*. Indeed, BG4 CUT&Tag performed in MCF7 cells revealed a distinct G4-peak in the promoter region of *IL17RA* ^56^, a gene that is only marginally expressed in this cell line. *IL17RA* encodes for the Interleukin 17 Receptor A, a proinflammatory cytokine secreted by activated T-lymphocytes and, therefore, not essential for breast cancer cell homeostasis. We generated sgRNA_IL17RA-20_ to target the IL17RA-G4 at 20 base pairs from its 5’ end and within a region that does not overlap with other regulatory elements of this promoter.

Following transfection with sgRNA_IL17RA-20_ or sgRNA_NTC_ and incubation with Cl-PDS_2_, we failed to detect any measurable changes in *IL17RA* expression levels (Fig. 5c). This observation indicates that targeting a G4 in a promoter of a transcriptionally inactive gene is not associated with gene-expression perturbation, linking tightly the functional relevance of G4s to the transcriptional levels of the genes associated. Considering that both G4s in the *HMGN1* and *IL17RA* promoters are equally detected in MCF7 by CUT&Tag ^56^ and targeted with similar sgRNA designs (within PAM sequence constraints), our findings suggest that the transcriptional levels linked to the targeted G4 can be used to anticipate the extent of transcriptional perturbation associated with ligand treatment. This model also explains the relatively modest transcriptional changes observed by the selective targeting of *c-MYC* and *PVT1* G4s, which are only moderately expressed in MCF7 cells. Increasing evidence in the literature suggests a model whereby G4s act as epigenetic factors to mark highly transcribed genes^12^. Our data support this model by showcasing how the extent of gene suppression/activation elicited by G4 ligands is linked to basal transcriptional levels, underscoring the relevance of maintaining G4 homeostasis in preserving transcriptional profiles characteristic of specific cell lines.

## Discussion

There is now substantial evidence to support that G4 structures form within endogenous chromatin and that their formation is intimately linked to transcriptional activity ^12^. For instance, G4s have been detected in the promoter regions of key oncogenes in cancer cells, and recent studies have demonstrated that they constitute critical structural features required to sustain high transcriptional rate, such as in the case of c-*MYC* ^20^. Over the past decades, the development of selective G4 ligands has revealed that stabilizing these structures often results in suppression of oncogene expression ^13^. Consequently, G4s have represented a therapeutically appealing target for decades. However, using G4-ligands for clinical application has not yet gained traction, reflecting two intrinsic limitations.

Firstly, the recognition mechanism leveraged by most G4 ligands relies on end-stacking interactions, which lack the selectivity to discriminate amongst different G4s - while effective at distinguishing G4s from duplex DNA. Given that the prevalence of G4s in the genome is highly cell-type specific, the lack of inter-G4 selectivity displayed by G4 ligands results in widespread transcriptional perturbations and inconsistent phenotypes across cell models used. Secondly, the binding affinity of individual ligands to different G4s varies broadly, making it difficult to pinpoint which G4s are functionally responsible for the biological responses elicited by any given ligand.

To address these limitations, tools enabling single-G4 targeting have been considered essential to unravel the fundamental biology regulated by specific G4s and to validate their therapeutic potential. Although several locus-directed and ligand-based strategies for single-G4 targeting have been reported^34,62–65^, intrinsic limitations associated with these methods summarized in Table S13, motivated us to develop ATENA-a CRISPR-guided platform in which catalytically inactive dCas9 is chemically functionalized with G4 ligands.

This system allows the positioning of a ligand in proximity to a specific G4 of interest using a short guiding RNA. We demonstrated that this approach enables transcriptional modulation attributable to G4 engagement at a single genomic locus

We initially optimized conditions for single-G4 targeting *in vitro*, before applying ATENA in cells to investigate the transcriptional role of a G4-structure located in the promoter region of the proto-oncogene *c-MYC*. While independent studies have previously reported *c-MYC* suppression upon treatment with G4-ligands^53^, it remains unclear whether this effect is exclusively mediated by engagement of ligands with the MYC-G4 or from broader transcriptomic changes induced by global G4-stabilisation. Indeed, Hurley and co-workers – who initially proposed that *c-MYC* downregulation was exclusively attributed to the targeting of MYC-G4^53^ - challenged the previous proposed model suggesting that transcriptional suppression was more likely a response to global G4-stabilization^18^.

Using ATENA, we observed that G4-mediated *c-MYC* suppression in MCF7 cells is minimal and limited to P1-controlled transcription. These findings are not only in line with what has been shown by recent genomic studies^20^ but were also fully recapitulated when using the MYC-G4 selective small molecule DC-34 ^24^. This strongly indicates that the biological response obtained by selective MYC-G4 targeting is limited to P1-mediated transcription, regardless of the targeting approach.

Importantly, we have noted that the MYC-G4 lies in proximity of key regulatory regions of the promoter, including the P1 promoter and the TATA-box sequence. Therefore, using CRISPR-based tools for selective MYC-G4 targeting can easily lead to misleading results when using sgRNAs targeting those regions and placing small molecules near these key regulatory elements. For instance, a recent study reported global *c-MYC* downregulation when using either dCas9-Nucleolin fusion or dCas9 poly-labeled with ten G4-ligands in tandem^34^. However, this effect was observed when targeting the same region as our sgRNA_MYC+58_, which places the dCas9 protein 9-bp apart from the TATA-box and we demonstrated to cause unspecific *c-MYC* downregulation (Fig. 2d). These findings indicated that placing ligands near core promoter elements can lead to transcriptional changes unrelated to G4-stabilization and that careful design of sgRNAs is required when using dCas9-based tools to avoid false positives. Additionally, our data suggested that the use of multiple ligands on a single dCas9 protein to induce transcriptional perturbation is likely leading to false positives by inducing local overcrowding at promoters ^34^. Similarly, using multiple sgRNAs on a single target can perturb the homeostasis of regulatory elements in a ligand-dependent manner, without necessarily reflecting G4-specific effects, and should thus be avoided ^34^.

Using the selective G4-antibody BG4, we confirmed that the P1-specific *c-MYC* downregulation induced by ATENA is accompanied by reduced protein accessibility at the MYC-G4 site, supporting the notion that G4 engagement underlies the observed transcriptional suppression by hampering G4-protein interactions. Notably, a similar effect was observed with DC-34, further suggesting that reduced BG4 binding, rather than the previously reported enhancement^34^, is to be expected when the ligand successfully engages the G4 within the *c-MYC* promoter. This also provides evidence that selective G4 targeting at the *c-MYC* promoter can be leveraged therapeutically to suppress its expression by interfering with protein-G4 interactions. However, this effect is limited to *c-MYC* transcription mediated by the P1 promoter and, therefore, heavily dependent on the cellular system investigated ^24^. For example, the substantial *c-MYC* downregulation, observed upon treatment with DC-34 in multiple myeloma cells^24^, is consistent with the high levels of P1-driven *c-MYC* expression characteristic of these cancers ^,67^, while it is ineffective in other cells that are less reliant on the P1 promoter for *c-MYC* expression (i.e., MCF7).

ATENA further enabled us to investigate the transcriptional response elicited when targeting the iM structure, also present within the *c-MYC* promoter. By decorating ATENA with an iM-selective peptide recently developed by the Waller’s group (RSV)^33^, we also observed transcriptional changes limited to the P1-promoter. However, when targeting the iM, we measured an increase of P1-driven transcription, rather than a reduction, suggesting that distinct DNA secondary structures might differently affect the expression at the same promoter. Moreover, this demonstrated the modularity of ATENA, which can be easily adapted to target any DNA structure of interest using the same design principles.

ATENA also enabled us to explore ligand-dependent variation in responses when targeting the same G4-structure. It is well established that G4-ligands can cause either transcriptional activation or repression depending on the G4-associated promoter, and this variability has often been attributed to context-dependent biological roles of the specific G4 investigated. However, this model contradicts genomic studies that strongly indicate a global association of G4-formation with transcriptional activation rather than a context-dependent function of these structures. Using ATENA in combination with CUT&Tag, we could demonstrate that the variation in gene expression responses at specific G4 sites stems not from inherent differences in G4 function, but rather from how structurally different ligands affect protein-G4 interactions.

We tested this by characterizing the different transcriptional responses observed upon targeting the G4 in the promoter of the long non-coding RNA *PVT1* when using two established G4 ligands: PDS and PhenDC3. The previously observed upregulation in *PVT1* transcription following PDS treatment^43^ was recapitulated with ATENA. This transcriptional increase was accompanied by increased BG4-binding at this G4-site, as quantified by BG4 CUT&Tag qPCR, suggesting that PDS binding might enhance local protein accessibility, acting as a molecular glue and, therefore, stimulate transcription. In contrast, PhenDC3 treatment resulted in transcriptional suppression - consistent with the more commonly reported effects of G4 ligands-demonstrating that structurally distinct ligands can drive divergent transcriptional responses at the same G4 site.

Notably, these ligand-dependent responses were also observed when using the free - not bound to dCas9 - PyPDS or PhenDC3, indicating that the lack of inter-G4 selectivity does not necessarily prevent these molecules from providing meaningful information on the specific G4 site. Taken together, these findings suggest that G4-ligands - while useful tools for perturbing G4-homeostasis and investigating the consequent biological responses - cannot be used to directly infer the native biological roles of G4s,but should instead be used to gain insights into the responses triggered by their binding to these structures. Moreover, our data strongly indicate that it is not safe to assume that different G4-ligands will lead to similar biological responses, which is a common assumption often reported in the literature.

Finally, we utilized ATENA to target uncharacterized G4s previously identified in MCF7 cells with CUT&Tag but never investigated their biological functions. Given that G4s are typically found at active promoters, we reasoned that a gene’s transcriptional status can strongly influence its responsiveness to G4-ligand targeting. We took advantage of ATENA to direct PyPDS to a G4-promoter of either a highly expressed gene (*HMGN1*) or a low-expressed one (*IL17RA*). Single targeting of the G4 located in the *HMGN1* promoter - unique to MCF7 cells and not detected in other cancer cell lines^56^ - led to the higher transcriptional suppression measured in MCF7 (a 99% reduction), confirming a strong link between transcriptional activity and ligand sensitivity that indicates a substantial contribution of the HMGN1-G4 in sustaining elevated expression levels of *HMGN1* in breast cancer cells.

We recently observed that PDS treatment in chemo-resistant ovarian cancer cells resensitizes them to chemotherapy, due to the enrichment of G4s in highly transcriptionally active genes that are essential to establish a chemo-resistant state, such as *WNT*^68^. Given the relevance of epigenetic changes in establishing dormancy in breast cancer^61^ and our observation linking expression levels of *HMGN1* with dormancy/awakening transition (Extended Data Fig. 5), targeting of the HMGN1-G4 with ligands might offer scope for the development of a therapeutic strategy to prevent awakening of dormant cells and cancer relapse.

Altogether, these results support a model in which the biological significance of G4 structures within the genome is contingent on the epigenetic landscape and the gene transcriptional status in a given cell line.

In conclusion, ATENA provides a robust method for targeting single G4s within the genome of living cells using distinct G4-ligands. This technology allowed us to dissect the diverse biological responses elicited by different ligands at the same G4 site, as well as the effects of targeting various G4s with the same ligand with high precision. We found that the chemical nature of the ligand used can perturb the local protein-binding homeostasis of promoter G4s differently, leading to either increased protein accessibility and transcriptional activation or decreased accessibility and transcriptional suppression. These effects can be monitored directly using BG4 CUT&Tag qPCR, providing an assessment of protein accessibility at the targeted G4 as a response to ligand binding. We demonstrated that some ligands could act as molecular glue of G4-protein interactions or as displacers, depending on the specific G4-targeted. This can be valuable for developing therapeutic agents based on G4-targeting, which can be tailored chemically to either diminish or amplify transcription based on their binding modalities. Moreover, ATENA enabled us to disentangle local effects triggered by individual G4-targeting from broader responses driven by global G4-stabilization obtained when using canonical G4-ligands. Interestingly, our data suggest that G4-ligands - despite their lack of inter-G4 selectivity - often recapitulate the transcriptional changes observed with selective targeting of G4s and can be used to infer the therapeutic potential of individual structures. Indeed, the response to G4-ligand treatment is mainly shaped by the intrinsic level of transcription of the targeted promoter, making the ligand responses highly epigenetic and cell-dependent.

We envisage that further development of ATENA can be leveraged to generate genome-wide screening of G4-functional response to various ligands in different cell lines. Additionally, this technology can be used to generate Halo-functional ligands and screen their ability to perturb protein accessibility at a given G4 site. We also anticipate that - given the minimal perturbation of ligands required for Halo-functionalization (chloroalkane) - ATENA is potentially a promising platform for gene-selective localization of other DNA-interacting therapeutics, including Topoisomerase inhibitors or cross-linking agents.

## Methods

Comprehensive synthetic protocols and purification methods for Cl-PDS_n_, Cl-PhenDC3_n_, and Cl-pep-RVS_n_ ligands are provided in the Supplementary Information (SI1).

### FRET-melting assay

Before each experiment, a solution containing 20 µM DNA ordered from Integrated DNA Technologies (IDT) (Table S1), labeled at 5’-with FAM and at 3’-with TAMRA, was freshly prepared in 10 mM lithium cacodylate buffer, pH 7.4 supplemented with 20 mM KCl. For the non-G4-forming sequence dsDNA26 (26 mer), the solution was supplemented with 150 mM KCl. The DNA solution was placed on an Eppendorf Thermomixer, annealed by heating at 95°C for 10 minutes, and then cooled at a rate of 0.5°C/minutes. The DNA was aliquoted into a 96-well RT-PCR plate (Thermo Fisher Scientific) and supplemented with increasing equivalent amounts of each ligand, resulting in a final oligonucleotide concentration of 0.4 µM. Fluorescence readings were performed on an Agilent Stratagene MX3000P, in the range of 25-95°C, in technical duplicate. The data points obtained were analyzed with GraphPad Prism 10, and the melting temperature was extrapolated from a Boltzmann sigmoidal function. Statistical significance was calculated using a two-tailed t-test in GraphPad Prism. p-value: ns > 0.05, * ≤0.05, ** ≤0.01, *** ≤0.001, **** ≤0.0001.

### CD-melting assay

Circular Dichroism (CD) melting experiments were conducted on an AVIV Biomedical Inc. (Lakewood, NJ, USA) Model 410 Circular Dichroism Spectrometer. Before each experiment, c-MYC-Pu22 (Table S1) was annealed at a concentration of 2 µM in lithium phosphate buffer (pH 7.4, 20 mM phosphate) supplemented with 1 mM KCl. When required, the ligand was added to the annealed solution and equilibrated for 1h before initiating the melting process.

Each experiment was performed in triplicate using a 10 mm optical path length quartz cuvette, with a 2 mm bandwidth. The temperature was ramped from 30°C to 95°C, with scans recorded every 5°C after a 2 min equilibration time. CD signals were measured at 264 nm (corresponding to the parallel G-quadruplex maximum).

The CD values (mdeg) at 264 nm were plotted as a function of temperature, normalized, and fit to a Boltzmann sigmoidal equation using GraphPad Prism 10.0. This analysis yielded the melting temperature (V50) of the G4 structure in the presence of increasing ligand equivalents. Statistical significance was calculated using a two-tailed t-test in GraphPad Prism. p-value: ns > 0.05, * ≤0.05, ** ≤0.01, *** ≤0.001, **** ≤0.0001.

### UV-binding assay

Oligonucleotides c-MYC-C52 and c-MYC-G52, purified by reverse-phase HPLC, were purchased from Eurogentec (Table S1). The oligonucleotide stock solutions were dissolved in MilliQ water and diluted to a final concentration of 250 µM in stock buffer (10 mM sodium cacodylate, pH 6.6) and annealed by heating at 95 °C for 5 min and cooling to room temperature overnight. The peptide was dissolved to 10 mM in DMSO, and 1 mM in DMSO was used in the experiment. UV-based binding curves were determined using the wavelength at 350 nm for 10 µM pep-RVS or 10 µM Cl-pep-RVS_4_ in stock buffer. DNA (250 µM in stock buffer) was added in increments up to 50 µM final concentration.

### dCas9-Halo purification

The expression vector for dCas9-Halo (Addgene #72269) was introduced into BL21 cells (D3, NEB-C2527H) by heat shock. Bacteria were then plated in LB agar plates and selected with Ampicillin (100 μg/ml). One bacteria colony was picked and grown overnight in a 50 ml flask at 37°C. Overnight precultures were diluted 1:100 into the main culture (LB media, Ampicillin 100 μg/ml) and incubated at 37°C, 200 rpm until an OD600 of 0.6. The expression of the recombinant dCas9-Halo was induced by using 1 mM of isopropyl β-D-1-thiogalactopyranoside (IPTG). Upon induction, the temperature was reduced to 16°C for dCas9-Halo expression. BL21-dCas9-Halo cells were harvested after overnight incubation at 16°C by centrifugation (3500 g, 20 minutes, 4°C). Cells were lysed in lysis buffer (50 mM sodium phosphate, 300 mM NaCl, pH 7.2, 20% glycerol, 0.1% Tween-20, 1 mM TCEP, 1 mg/ml lysozyme, 1 mM PMSF, 1 mM MgCl_2,_ and 5U/g of culture Benzonase). Cell lysates were then homogenized, and cell debris was removed by centrifugation (16000xg, 15 minutes, 4°C). The soluble fraction of the lysate was filtered and then loaded onto a cobalt column, and the bound protein was eluted with 150 mM imidazole. After imidazole removal and concentration by using a 50,000-MWCO centrifugal filter (Millipore Amicon), fractions were loaded into a 1-ml HiTrap SP XL (GE Healthcare), ÄKTA pure 25 for cationic exchange chromatography, and elution was performed with a gradient from 0% to 70% of Buffer (50 mM HEPES pH 7.5, 1 M KCl, 1mM TCEP). Purest fractions were pooled and then used for gel filtration chromatography. Gel filtration was performed using a Superdex 200 10/300 GL column, and isocratic elution was performed in a buffer containing 50 mM HEPES (pH 7.5), 150 mM KCl, and 1 mM TCEP. Protein aliquots in storing buffer (50 mM HEPES, pH 7.5, 150 mM KCl, 20% glycerol) were snap-frozen and stored at -80°C.

### BG4 purification

BL21 *E. coli* competent cells (D3, NEB-C2527H) were transformed with a pSANG10-3F-BG4 plasmid (Addgene #55756). Single colonies were inoculated in 2xYT medium supplemented with 2% glucose and 50 μg/ml kanamycin and incubated at 30 °C at 250 rpm overnight. The starting culture was diluted 1:500 into ZYM-5052 autoinduction medium prepared following the previously described method^69^ supplemented with 50 μg/ml kanamycin. The culture was then incubated at 37 °C for 6 hours at 250 rpm, followed by an overnight incubation at 25 °C at 280 rpm. The culture was centrifuged at 4,000 x *g*, 4 °C for 30 minutes, and the collected pellet lysed in 80 ml ice-cold TES buffer (50 mM Tris-HCl pH 8.0, 1 mM EDTA pH 8.0, 20% sucrose) and incubated on ice for 10 minutes. This was followed by 15 minutes of incubation in ice-cold TES (diluted 1:5), supplemented with 15 U/mL benzonase, 2 mM MgSO4, and a protease inhibitor cocktail. The lysate was centrifuged at 16,000 x g for 20 minutes. The supernatant was collected and incubated with Nickel resin (Sigma, P6611), previously equilibrated in PBS, at 4 °C for 1 hour in rotation. The complex was washed three times with wash buffer (PBS, 100 mM NaCl, 10 mM imidazole). Then, BG4 was eluted in 4 mL of elution buffer (PBS, pH 8.0, 250 mM imidazole) and dialyzed against PBS at 4 °C using GeBaflex tubes (Generon). Purified BG4 was snap-frozen in liquid nitrogen and stored at -80 °C.

### dCas9-binding competition assay

dCas9-Halo purified protein (4 µM) was incubated with several dilutions of Cl-PDS_n_ or Cl-PhenDC3_n_ probes or with commercially available Halo-TAMRA for the positive control (Promega, G8252) for 30 minutes at room temperature in PBS. Pretreated dCas9-Halo proteins were then incubated with 5 μM Halo-TAMRA, or Cl-PDS_n_ for the positive control, and samples were loaded in an SDS-PAGE gel (8% polyacrylamide) and run at 180V for 45 minutes. Gels were then imaged using a Typhoon FLA 9500 (GE) with the TAMRA filter set (excitation at 542 nm) and then stained with Coomassie. Images of Coomassie-stained gels were then acquired with ImageQuant LAS 4000 (Cytiva).

Each band intensity was then quantified using Image Studio software and normalized for the corresponding Coomassie signal, expressed as a percentage (with 100% labeling corresponding to the positive control). IV-CP_50_ values were determined using nonlinear regression (dose-response inhibition curves with constrained fitting) in GraphPad Prism with n = (number of independent experiments) = 2.

### In vitro transcription for sgRNA synthesis

Oligonucleotides containing sgRNA sequences (Table S2) were ordered from Integrated DNA Technologies (IDT), and a PCR was set up to generate the corresponding duplex following the previously described method^70^. Following the manufacturer’s instructions, the DNA product was transcribed using the HiScribe T7 High Yield RNA Synthesis Kit (NEB, E2040S). The resulting sgRNA was purified using the RNeasy Mini kit (74104) and stored at -80°C.

### dCas9-Halo_FRET assay

Before the experiment, Cy5-3’ end-labeled and Cy3-5’ end-labeled oligos (Table S1) were annealed in 10 mM Tris HCl pH 7.5, 100 mM KCl at a final concentration of 1 μM (95 °C for 10 minutes and left for overnight cooling down at room temperature). dCas9-Halo purified protein at a final concentration of 4 μM was incubated with Cl-PDS_n_ probes in a 1:2.5 ratio for 45 minutes in binding buffer (20 mM HEPES, pH 7.5, 100 mM KCl, 1 mM MgCl_2_, 1 mM TCEP, 10% glycerol). The pretreated dCas9-Halo was then incubated with 120 pmol of purified sgRNA for 20 minutes at room temperature, followed by incubation with 200 nM of pre-annealed oligos at 37°C for 1h. The complexes were then loaded into an 8% polyacrylamide native gel and ran for 120 minutes at 4°C in 1x TBE supplemented with 2 mM MgCl2. Gels were then imaged using a Typhoon FLA 9500 (GE) with the Cy3 and Cy5 filter sets. Analysis of the fluorescent band intensity (I) and relative FRET efficiency (E) was performed using a Python code. Particularly, FRET efficiency was calculated as: E=I_Cy5_ /(I_Cy3_ +I_Cy5_). The signals in both channels were normalized for the background and the sgRNA NTC control, and for every sgRNA, the ΔFRET is defined as: (E+ligand) - (E−ligand). Statistical significance was calculated using a two-tailed t-test in GraphPad Prism; p-value: ns > 0.05, * ≤0.05, ** ≤0.01, *** ≤0.001, **** ≤0.0001 with n = (number of independent experiments) = 2.

### Cloning of lentiviral dCas9-Halo construct

The lentiviral dCas9-Halo plasmid was cloned using Gibson assembly. A lentiviral backbone containing already dCas9 (Addgene #61425) was digested using BamHI and BsrGI (New England Biolabs, R3575S, R3136S) and then assembled with the HaloTag sequence amplified (Table.S3) from pET302-6His-dCas9-Halo (Addgene #72269) in a 1:3 molar ratio of backbone: insert using HiFi DNA Assembly Mix (E2621S, New England Bioscience) following manufacturing protocols. Post incubation, assembled products were diluted with water, and 5 μL of the product was transformed by heat shock into 10-beta competent cells (New England Biolabs, C3019I). Cells were then plated on agarose plates (supplemented with ampicillin 100 μg/mL) for overnight outgrowth at 37 ℃. Single clones were picked and grown in 5 mL of Amp LB media overnight at 37 °C while shaking. Plasmid DNA was purified from cells using the Promega PureYield plasmid miniprep system (Promega, A1223) and sequenced using the Genewiz service.

The positive plasmid was further modified by substituting the blasticidine resistance gene with the mCherry coding sequence using Gibson assembly after digestion with BsrGI and EcoRI enzymes (Table S3). Positive clones were then sequenced using the Genewiz service.

### Cloning of sgRNA in pLG1 plasmid

The pLG1 backbone (Addgene #109003) was digested with BstXI and BlpI (FastDigest, ThermoFisher) for 1 hour at 37°C. Oligos containing the sgRNA sequence (Table S4) were ordered from IDT and annealed in water at 10 μM in a thermocycler (37°C, 30 minutes-95°C, 5 minutes and ramp down 5 degrees/minute to 25°C). The annealed oligos were diluted (1:50) and then assembled with the digested plasmid in a 1:2 molar ratio of backbone: insert using HiFi DNA Assembly Mix (E2621S, New England Bioscience) following the manufacturer’s protocols. Post-incubation, the assembled products were diluted 1:2 in water, and 5 μL of the product was transformed by heat shock into 10-beta competent cells (New England Biolabs, C3019I). Cells were then plated on agarose plates (supplemented with 100 μg/mL ampicillin) for overnight outgrowth at 37 ℃. Single clones were picked and grown overnight in 5 mL of Amp LB medium at 37 °C in a rotating shaker. Plasmid DNA was purified from cells using the Promega PureYield plasmid miniprep system (Promega, A1223) and sequenced using the Genewiz service.

### Mammalian cell culture

HEK293FT cells were cultured in DMEM GlutaMAX™ (Thermo Fisher Scientific, 10566016) supplemented with 1x penicillin-streptomycin (Thermo Fisher Scientific, 15140122) and 10% (vol/vol) fetal bovine serum (FBS) and maintained at 37 ℃ and 5% CO2.

MCF7 cells were cultured in DMEM GlutaMAX™ (Thermo Fisher Scientific, 10566016) supplemented with 1x β-Estradiol (Sigma, 50-28-2), 1x penicillin-streptomycin (Thermo Fisher Scientific, 15140122), and 10% (vol/vol) fetal bovine serum (FBS) and maintained at 37 ℃ and 5% CO2.

### Lentivirus production and transduction

For lentiviral production, HEK293T cells were seeded at 3.8×10^6^ in a 10 cm tissue culture plate the day before transfection. 8.4 μg of the envelope plasmids pCMV-VSV-G (Addgene #8454) and 6.4 ug of packaging plasmid R8.74 (Addgene #22036) were co-transfected along with 2.1 μg of the target plasmid (dCas9-Halo-T2A-mCherry) using polyethyleneimine (PEI-Linear, MW 25000) in a μg DNA: ug PEI ratio of 1:3. Viral supernatant was harvested after 48h and 72h, and before usage, it was filtered using a 0.45 μm filter unit. MCF7 cells were plated on 6-well plates the day before infection. They were infected with lentiviruses in DMEM 10% FBS, β-Estradiol, in the presence of polybrene with a final concentration of 10 μg/mL. Half of the media was changed the day after, and cells were grown for one week and examined by flow cytometry (Attune NxT, ThermoFisher) to confirm successful transduction (YL2-Channel). After genotyping (Table S3), cells were single-cell sorted using FACS BD FACS Diva 9.0.1 (Fig.SI-3).

### Western blot

MCF7 stably expressing dCas9-Halo cells were harvested and processed using 1x RIPA buffer (Merck,20-188) supplemented with 1x Protein inhibitor cocktail. The total protein concentration of the cleared lysate was then measured by Bradford assay (Pierce™ Bradford Plus Protein Assay Kits, Thermo Fisher Scientific, 23236) at 595 nm using a ClarioStar plate reader. A total of 20/25 µg protein was then loaded into an 8% polyacrylamide SDS-PAGE gel and transferred to a pre-assembled PVDF membrane (Trans-Blot Turbo Mini 0.2 µm PVDF Transfer Packs, Bio-Rad) using a semi-dry method (Bio-Rad). The membrane was blocked (5% milk in TBS-T) and incubated with primary antibodies (1:1000 Cas9 antibody-14697T, 1:1000 GAPDH antibody-2118T, Cell Signaling Technology) overnight at 4 °C. The membrane was then washed in TBS-T and incubated with secondary antibodies (1:10,000) goat anti-rabbit/mouse HRP-Advansta, R-05071-500, R-05072-500) for 1h at room temperature. After three washing steps of the membrane in TBS-T, 0.5 ml of HRP substrate (Merck, WBLUC0100) was added to the top of the membrane, and the excess was removed. The signal was developed using Image Quant LAS 4000 (Cytiva) with the following settings: chemiluminescence for the WB signal and Cy5 for visualizing the protein marker.

### CAPA assay

MCF7 cells stably expressing dCas9-Halo were plated the day before at 30×10^3 cells/ well in 96-well plates pre-coated with Poly-D-Lysine (Thermo Fisher Scientific, A3890401). Cells were incubated for 2h in the presence of serial dilutions of Cl-PDS_n_, Cl-PhenDC3_n_,or Cl-pep-RVS_n_ probes ranging from 0.2μM to 10 μM (5% CO2, 37 °C). Cells were then washed two times with media (every washing step included 10 minutes of incubation, 5% CO2, 37 °C) and incubated with HaloTag® Oregon Green® Ligand (Promega, G2801) for 45 minutes, followed by two washing steps (10 minutes,5% CO2, 37 °C). Lastly, cells were washed with DPBS, treated with 0.25% trypsin, and resuspended in FACS buffer (5% BSA in DPBS) before undergoing flow cytometry analysis (Attune Nxt). Data analysis was performed in FlowJo 10.9.0. Mean fluorescence values from two biological replicates (each with three technical replicates) were normalized to the positive-control signal and expressed as percent labeling. Following this, CP_50_ values were obtained by nonlinear regression (dose–response inhibition curves with constrained fitting) in GraphPad Prism (n = 2).

### Transfection and incubation with G4 ligands

MCF7 cells stably expressing dCas9-Halo were plated the day before at 25×10^3 cells/ well in 96-well plates pre-coated with Poly-D-Lysine (Thermo Fisher Scientific, A3890401). According to the manufacturer’s protocol, 50 ng of guide expressing plasmid was transfected using Lipofectamine 3000 (Thermo Fisher Scientific, L3000001). The day after, cells were incubated with either DMSO (0.5%) or Cl-PDS_n,_Cl-PhenDC3_n,_ or Cl-pep-RVS_n_ probes and incubated for 48h.

### RNA isolation

RNA was harvested 72 hours post-transfection. Cells were washed with 100 μL of 1X DPBS (Gibco, 14190144) and incubated with RLT buffer from RNeasy mini kits (Qiagen, 74104) according to the manufacturer’s instructions. Following the manufacturer’s instructions, the eluted RNA was reverse-transcribed into cDNA using the RevertAid cDNA Synthesis Kit (K1621, Thermo Fisher Scientific).

### Quantitative Real-Time PCR (RT-qPCR)

RT-qPCR reactions were prepared as follows: cDNA was mixed with 500 nM of forward and reverse primer (Table S5) and 5μL of Fast SYBR Green Master Mix (Applied Biosystems: 4385612) in a final volume of 10 μL. RT-qPCR was performed on a 96-well plate format using an Agilent Stratagene Mx3000P machine with the following program: 95 °C for 20 seconds, 40× (95 °C for 3 s, 60 °C for 30 s), 95 °C for 15 s, 60 °C for 1 minute, and 95 °C for 15 s (melting curve). The C_t_ values obtained were used to calculate the relative fold-change in gene expression using the ΔΔC_t_ method and normalized to the housekeeping gene *GAPDH* in both treated and untreated samples. Statistical significance was calculated using a Welch-corrected two-tailed t-test in GraphPad Prism; p-value: ns > 0.05, * ≤0.05, ** ≤0.01, *** ≤0.001, **** ≤0.0001.

### BG4 CUT&Tag

Cells were plated in a 6-well plate format one day before transfection, following the previously described transfection protocol. 24h after transfection, cells were incubated with Cl-PDS_n_ probes for 6h and then harvested, counted, and inspected for viability. To prepare nuclei, 200k cells per reaction (plus 10% excess) were centrifuged for 3 minutes at 600×*g* at room temperature and then resuspended in nuclei extraction buffer (20 mM HEPES–KOH, pH 7.5, 10 mM KCl, 0.1% Triton X-100, 20 mM Glycerol, 0.5 mM Spermidine, 1X Roche cOmplete Mini EDTA-free Protease Inhibitor (Roche, 11836170001) and incubated for 10 minutes on ice. Nuclei were spun at 600 x g for 3 min at 4 °C and then gently resuspended in 200 μL/per reaction cold Nuclei Extraction Buffer. A 2 μL aliquot was taken and mixed with 2 μL trypan blue to examine nuclei integrity under the microscope. In the meantime, ConA beads (BioMag®Plus Concanavalin A, BP531) were prewashed and equilibrated in binding buffer (20 mM HEPES–KOH, pH 7.5, 10 mM KCl, 1 mM CaCl_2_, 1 mM MnCl_2_). Nuclei for each condition that needed to be tested were mixed with 10 μL of beads and incubated for 10 minutes at RT while shaking. After checking the binding of the beads to the ConA beads under the microscope (2 μL sample + 2 μL trypan blue), the samples were washed two times with 300 μL of 1%BSA antibody buffer (20 mM HEPES–KOH, pH 7.5, 150 mM KCl,0.5 mM Spermidine, 1X Roche cOmplete Mini EDTA-free Protease Inhibitor, 5% Digitonin, 2mM EDTA, and 1%BSA). The tubes were placed on the magnetic rack, and the liquid was withdrawn. Samples were then resuspended in 50μL/ reaction of 1%BSA antibody buffer and incubated for 1h at RT while shaking. The samples (3 technical replicates for each condition) were incubated with 216 nM of an in-house prepared BG4 antibody and 0.5 μL of H3K27me3 (Cell Signaling Technology,9733S) antibody for the positive control (1 replicate for each condition) and left overnight at 4°C while shaking.

The day after, samples were checked to ensure that they were bound to the beads and then washed two times with Dig-wash buffer (20 mM HEPES–KOH, pH 7.5, 150 mM KCl,0.5 mM Spermidine, 1X Roche cOmplete Mini EDTA-free Protease Inhibitor, 5% Digitonin). Samples containing BG4 and samples for the negative control were resuspended in 50 μL of Dig-wash buffer containing 2 μL of anti-flag antibody (DYKDDDDK Tag Antibody, 2368S) and incubated for 1h at RT while shaking. Then, the BG4 and negative samples were washed three times in 100 μL Dig wash buffer, resuspended in 50 μL Dig wash buffer supplemented with 0.5 μL of tertiary antibody (anti-rabbit, Abcam, ABIN101961) and incubated for 1h at RT while shaking. Samples were then washed three times with 100μL Dig-wash buffer, resuspend in 50 µL Dig-300 buffer (20 mM HEPES–KOH, pH 7.5, 300 mM KCl, 0.5 mM Spermidine, 0.01% Digitonin, 1X Roche cOmplete Mini EDTA-free Protease Inhibitor) containing pA-Tn5 (CUTANA™ pAG-Tn5 for CUT&Tag, 15-1017) adapter complex (1:20) and incubated for 1h at RT while shaking. Samples were then washed three times in Dig-300 buffer, resuspended in 100 μL of Tagmentation buffer (Dig-300 buffer supplemented with 10 mM MgCl_2_) and incubated for 1h at 37°C. Samples were then, washed twice with TAPS buffer (10 mM TAPS buffer-J63268.AE, Thermo Scientific Chemicals, 0.2 mM EDTA), mixed by vortexing in 100 μL of Protenaise K buffer (0.5mg/mL Proteinase K, Thermo Fisher Scientific EO0492/EO0491, 0.5% SDS in 10mM Tris-HCl buffer pH 8.0) and incubate 1 hr at 55 °C. Samples were then purified with Zymo DNA Clean & Concentrator-5 (D4013) and samples were then, amplified using NEBNext HiFi 2x PCR Master mix (NEB, M0541) with the following set up (Initial 72 °C for 5 minutes, 20X cycle (98 °C for 30 sec, 98 °C for 10 sec, 63 °C for 10 sec) and 72°C for 1 minutes). Samples were then purified using Ampure XP beads (A63880) according to the manufacturer’s instructions. Quality control of the sample was performed using TapeStation ScreenTape HSD1000 (5067-5584) and TapeStation ScreenTape HSD1000 reagents (5067-5585) according to the manufacturer’s instructions on a TapeStation (TapeStation Agilent 4150).

### BG4 CUT&Tag-qPCR

CUT&Tag libraries were diluted 1:10, and 2 µL of the diluted CUT&Tag library was mixed with 1 μM of primer mix (Table S6) and 5 μL of SYBR Green PCR Master Mix (Applied Biosystems: 4385612) according to the following protocol 20 seconds at 95 °C, 39 cycles of 10 seconds at 95 °C, 30 seconds at 60 °C, and 10 seconds at 72 °C, and finally heating to 90 °C. The C_t_ values obtained were used to calculate the relative fold-change in gene expression using the ΔΔC_t_ method and assess the relative fold change at G4 target sites when compared against G4-positive regions (MAZ, RBBP4, RPA3). Statistical significance was calculated using a Welch-corrected two-tailed t-test in GraphPad Prism; p-value: ns > 0.05, * ≤0.05, ** ≤0.01, *** ≤0.001, **** ≤0.0001.

### Histone modifications CUT&Tag

The CUT&Tag assay was performed using the EpiCypher CUTANA Direct-to-PCR CUT&Tag Protocol v1.7, which included all the mentioned materials and buffer recipes with minor modifications. Briefly, cryopreserved nuclei were thawed quickly and immobilized to Concanavalin A (ConA) Conjugated Paramagnetic Beads (Epycypher, cat. 21-1401). Bead-bound nuclei were spiked with 2 µL of spike-in SNAP-CUTANA™ K-MetStat Panel (1:50, Epicypher, cat. 19-1002) and immediately resuspended and incubated in Primary Antibody diluted in Antibody buffer at the manufacturer’s CUT&Tag recommended dilutions, at 4°C by nutation overnight. The next day, the primary antibody was removed, and nuclei were incubated with either anti-Rabbit or anti-Mouse Secondary Antibodies at Room Temperature (RT) in Digitonin 150 buffer for 1h with nutation. Nuclei were washed twice with wash 150 buffer and incubated with CUTANA pAG-Tn5 enzyme (Epicypher, cat. 15-1017) diluted in Digitonin 300 buffer for 1h at RT with nutation and thoroughly rewashed twice. Nuclei were resuspended in Tagmentation Buffer containing MgCl2 and incubated 1 h at 37°C.

The reactions were then washed with Post-tagmentation buffer and incubated in SDS Release Buffer at 58°C for 1h to quench the tagmentation reaction. Then, SDS Quench Buffer is added to neutralize SDS, as well as NEBNext High-Fidelity 2 × PCR Master Mix (NEB, cat. M0541L), and 2.5 µM of universal i5 primer and unique i7 primer sequences previously published in Buenrostro et al. 2015 (IDT technologies). Libraries were amplified at 14 cycles for abundant histone modification samples (H3K27me3), and 16 cycles for less abundant marks (H3K4me3) and negative control IgG following the cycling parameters outlined in the EpiCypher® CUTANA™ Direct-to-PCR CUT&Tag Protocol v1.7. Library clean-up was performed using 1.3x SPRI beads (Beckman Coulter, cat. B23319) to recover ∼75 bp DNA fragments. The beads-DNA were incubated at RT for 5 minutes, followed by two washes with 85% ethanol. Libraries were eluted in 15 μL of 0.1x TE buffer, quantified using Qubit™ fluorometer per manufacturer’s instructions, and quality controlled for fragment length enrichment using the Agilent TapeStation Bioanalyzer® with High Sensitivity D1000 reagents. The following antibodies were used for CUT&Tag: H3K4me3 (Epycypher, cat 13-0041, 1:50), H3K27me3 (Cell signalling, cat 9733T, 1:50), IgG (Epycypher, cat 13-0042, 1:1000), Anti-Rabbit Secondary Antibody (Epycypher, cat 13-0047, 1:100) and Anti-Mouse Secondary Antibody (Epycypher, cat 13-0048, 1:100). Libraries were sequenced at 10 million reads per sample on Paired-End platform of 150bp per fragment (PE150) by our sequencing provider, Novogene, on a NovaseqX platform from Illumina.

CUT&Tag data was processed on the Imperial College London HPC using an adapted version of the pipeline developed by Dr Ye Xheng from the Steven Henikoff lab, available on https://github.com/clabanillas/cutnTag_processing. This pipeline includes quality control, adapter trimming, alignment, filtering, and peak calling. Peaks for H3K27me3 were called using SEACR with no built-in normalisation, against control IgG in stringent mode. Peaks for H3K4me3 were called with IgG control with MACS2 callpeak command (--keep-dup all --nomodel --shift -75 --extsize 150 -q 0.01).

### ATAC-seq

scATAC-seq was performed on a Chromium platform (10x Genomics) using Chromium Single Cell ATAC Reagent Kit’ V1 chemistry (manual version CG000168 Rev C) and Nuclei Isolation for Single Cell ATAC Sequencing (manual version CG000169 Rev B) protocols. Nuclei suspensions were prepared to get 10,000 nuclei as target nuclei for recovery. Final libraries were loaded onto a Novaseq 6000 platform (Illumina) to obtain 50,000 reads per nucleus with a read length of 2 × 50 bp.

A pseudobulk of single-cell sequencing readout was prepared by concatenating all reads containing the transposase adaptor sequence barcode per sample. Data processing was done according to the following repository: https://github.com/harvardinformatics/ATAC-seq. Briefly, raw paired-end FASTQ files were subjected to quality control using FastQC, and adapter trimming was performed with FASTP (--detect_adapter_for_pe -l 20). Reads were then aligned to the human genome (T2T-CHM13v2.0) using Bowtie2 with parameters optimized for ATACseq (--very-sensitive -X 700).

Mitochondrial reads were removed with Samtools. Duplicates were removed using PICARD (picard-2.27.4-0/picard.jar). Then multimapped reads were removed with samtools (-h -q 30), as well as unmapped, mate unmapped, not primary alignment, reads failing platform, duplicates (-h -b -F 1804), and properly paired reads -f 2 were retained. Reads aligned to blacklisted regions were removed.

Coordinates were shifted with deeptools alignmentSieve command (--numberOfProcessors max – ATACshift). Normalised genome coverage tracks were generated with the BEDtools genomecov function (--numberOfProcessors max --binSize 10 --normalizeUsing RPGC --effectiveGenomeSize 2786136059) using the effective genome size for T2T. Peaks were called using the MACS2 callpeak command (-q 0.01 --keep-dup all).

### RNAseq sample preparation and bioinformatic analysis

For transcriptome-wide gene expression changes in transfected samples, cells were plated the day before in 6-well plates, and 24h after transfection, they were incubated with either 0.5% DMSO or 2.5 μM Cl-PDS_2_ for 6 hours. For DC-34/PyPDS-treated samples, cells were plated in a 6-well plate the day before treatment with 5 μM and 2.5 μM of the compound for 6h. Cells were harvested and washed with 1x DPBS (300xg, 5 minutes). RNA was then extracted using RNeasy mini kits (Qiagen, 74104) according to the manufacturer’s instructions. The quality of the extracted RNA was assessed by TapeStation (High Sensitivity RNA Screen Tape, Agilent) and sent to Novogene service for sequencing. RNA libraries were prepared using Novogene kit for library preparation and then sequenced with Illumina NovaSeq X-plus. CASAVA (version 1.8) was used to perform base calling and Phred score. The quality of the reads was assessed with FastQC (Fastp v.0.23.1). Sequence alignment to the reference genome (GRCh38-NCBI:GCA_000001405.27) was performed using HISAT2. Sorted bam files were generated with (Hisat2 v.2.0.5), and gene counts using FPKM.

Raw count data were then imported into DESeq2 (v1.40.2), and differential gene expression analysis was conducted with three comparisons: (1) the generic drug effect in control (NTC) cells treated with the ligand versus mock (DMSO), (2) the effects of the sgRNA (sgRNA_MYC-19_) under mock conditions and data set (2) filtered for NTC. DEGs were identified based on adjusted p-values (FDR < 0.05) and log2 fold-change thresholds (|log2FC| ≥ 1).

### Cell proliferation assay

MCF7 cells, wild-type or stably expressing dCas9-Halo were plated one day before the experiment at 24 x 10^3 cells/well in a 96-well plate. Cells were then treated 24h later with serial dilutions (ranging from 0.25 μM to 10 μM) of Cl-PDS_2_, Cl-PhenDC3_2_, Cl-PDC_2_, PyPDS, PhenDC3, PDC, DC-34 or Cl-pep-RVS<u>_n_</u> and 0.5% DMSO as a negative control. After 48h, cells were incubated with 20 µL/well of CellTiter96^®^AQ_ueous_One Solution Reagent for 2 h (5% CO_2_, 37 °C), before acquiring the absorbance value at 490 nm using a ClarioStar plate reader. The data were fitted to a sigmoidal dose-normalized response curve with a variable slope. Statistical analysis of curve-fit parameters was performed by independently fitting data from separate biological experiments, followed by comparison of the resulting curve-fit parameters using Extra sum-of-squares F test in GraphPad Prism. p-value: ns > 0.05, * ≤0.05, ** ≤0.01, *** ≤0.001, **** ≤0.0001.

### Data availability

All new datasets have been deposited in the gene expression omnibus (GEO): GSE279769. To review GEO accession GSE279769 Enter token “itwzseukjhwfjon” into the box: https://www.ncbi.nlm.nih.gov/geo/query/acc.cgi?acc=GSE279769

## Acknowledgments

M.D.A is supported by a Biotechnology and Biological Sciences Research Council (BBSRC) David Phillips Fellowship (BB/R011605/1) and is a Lister Institute Research Prize holder (2022). S.P.N. acknowledges support from the Engineering and Physical Sciences Research Council (EPSRC) (EP/S023518/1). T.E.M acknowledges support from the Engineering and Physical Sciences Research Council (EPSRC) (EP/S023518/1). J.S.S. is supported by the Intramural Research Programs of the National Institutes of Health, National Cancer Institute (NCI), Center for Cancer Research, Project BC011585. D.G. was supported by a BBSRC grant (BB/W001616/1). This work was supported by a Leverhulme Trust Research Grant (RPG-2022-184). E.C. was supported by FWO (12B1923N). We are thankful to Prof Luca Magnani for providing MCF7 cells and for helpful discussions. The authors acknowledge Prof Ramon Vilar, Dr Michele Stasi, and Dr Koustav Pal for insightful discussions. The authors also acknowledge Dr Denise Liano, Dr Aisling Minard, Dr Souroprobho Chowdhury and Dr Anna Di Porzio for initial experimental efforts on this project.

## Authors Contribution

M.D.A. conceived and supervised the research project. S.P.N. designed the research with M.D.A. and performed all the experiments unless otherwise specified. E.C, L.S.L., R.N., A.J.L., E.F., M.Z., T.E.M. have contributed to synthesizing the G4-ligands. E.C. and A.J.L. perform FRET melting assays. E.C. performed the synthesis and chemical characterization of the pep-RVSn ligands and CD melting experiments. S.G. performed with S.P.N. BG4 CUT&Tag experiments. C.S.C. and L.M. performed and analyzed ATAC-seq and CUT&Tag of histone marks. Z.W. and D.G. advised on i-motif-peptides and D.G. performed binding studies on the pep-RVSn ligands. G.F. helped with chromatin prep. J.S.S. conceived DC-34 and helped edit the manuscript. C.R.F synthesized DC-34 used in this study. M.D.A. wrote the manuscript supported by S.P.N with input from all authors. All authors contributed to critical discussion and data interpretation.

## References

1 Davis, J. T. G-quartets 40 years later: from 5’-GMP to molecular biology and supramolecular chemistry. Angew. Chem. Int. Ed. Engl. 43, 668–698 (2004).

2 Gellert, M., Lipsett, M. N. & Davies, D. R. Helix formation by guanylic acid. Proc. Natl. Acad. Sci. U. S. A. 48, 2013–2018 (1962).

3 Parkinson, G. N., Lee, M. P. & Neidle, S. Crystal structure of parallel quadruplexes from human telomeric DNA. Nature 417, 876–880 (2002).

4 Biffi, G., Tannahill, D., McCafferty, J. & Balasubramanian, S. Quantitative visualization of DNA G-quadruplex structures in human cells. Nat. Chem. 5, 182–186 (2013).

5 Di Antonio, M. et al. Single-molecule visualization of DNA G-quadruplex formation in live cells. Nat. Chem. 12, 832–837 (2020).

6 Summers, P. A. et al. Visualising G-quadruplex DNA dynamics in live cells by fluorescence lifetime imaging microscopy. Nat. Commun. 12, 162 (2021).

7 Lyu, J., Shao, R., Kwong Yung, P. Y. & Elsasser, S. J. Genome-wide mapping of G-quadruplex structures with CUT&Tag. Nucleic Acids Res. 50, e13 (2022).

8 Esnault, C. et al. G4access identifies G-quadruplexes and their associations with open chromatin and imprinting control regions. Nat. Genet. 55, 1359–1369 (2023).

9 Hansel-Hertsch, R. et al. G-quadruplex structures mark human regulatory chromatin. Nat. Genet. 48, 1267–1272 (2016).

10 Hansel-Hertsch, R. et al. Landscape of G-quadruplex DNA structural regions in breast cancer. Nat. Genet. 52, 878–883 (2020).

11 Chambers, V. S. et al. High-throughput sequencing of DNA G-quadruplex structures in the human genome. Nat. Biotechnol. 33, 877–881 (2015).

12 Robinson, J., Raguseo, F., Nuccio, S. P., Liano, D. & Di Antonio, M. DNA G-quadruplex structures: more than simple roadblocks to transcription? Nucleic Acids Res. 49, 8419–8431 (2021).

13 Balasubramanian, S., Hurley, L. H. & Neidle, S. Targeting G-quadruplexes in gene promoters: a novel anticancer strategy? Nat. Rev. Drug Discov. 10, 261–275 (2011).

14 Kosiol, N., Juranek, S., Brossart, P., Heine, A. & Paeschke, K. G-quadruplexes: a promising target for cancer therapy. Mol. Cancer 20, 40 (2021).

15 Marsico, G. et al. Whole genome experimental maps of DNA G-quadruplexes in multiple species. Nucleic Acids Res. 47, 3862–3874 (2019).

16 Lago, S. et al. Promoter G-quadruplexes and transcription factors cooperate to shape the cell type-specific transcriptome. Nat. Commun. 12, 3885 (2021).

17 Figueiredo, J., Mergny, J. L. & Cruz, C. G-quadruplex ligands in cancer therapy: Progress, challenges, and clinical perspectives. Life Sci. 340, 122481 (2024).

18 Boddupally, P. V. et al. Anticancer activity and cellular repression of c-MYC by the G-quadruplex-stabilizing 11-piperazinylquindoline is not dependent on direct targeting of the G-quadruplex in the c-MYC promoter. J. Med. Chem. 55, 6076–6086 (2012).

19 Galli, S., Flint, G., Ruzickova, L. & Di Antonio, M. Genome-wide mapping of G-quadruplex DNA: a step-by-step guide to select the most effective method. RSC Chem. Biol. 5, 426–438 (2024).

20 Esain-Garcia, I. et al. G-quadruplex DNA structure is a positive regulator of MYC transcription. Proc. Natl. Acad. Sci. U. S. A. 121, e2320240121 (2024).

21 Spiegel, J. et al. G-quadruplexes are transcription factor binding hubs in human chromatin. Genome Biol. 22, 117 (2021).

22 Zanin, I. et al. Genome-wide mapping of i-motifs reveals their association with transcription regulation in live human cells. Nucleic Acids Res. 51, 8309–8321 (2023).

23 Zeraati, M. et al. I-motif DNA structures are formed in the nuclei of human cells. Nat. Chem. 10, 631–637 (2018).

24 Calabrese, D. R. et al. Chemical and structural studies provide a mechanistic basis for recognition of the MYC G-quadruplex. Nat. Commun. 9, 4229 (2018).

25 Cadoni, E., De Paepe, L., Manicardi, A. & Madder, A. Beyond small molecules: targeting G-quadruplex structures with oligonucleotides and their analogues. Nucleic Acids Res. 49, 6638–6659 (2021).

26 Minard, A. et al. A short peptide that preferentially binds c-MYC G-quadruplex DNA. Chem. Commun. (Camb) 56, 8940–8943 (2020).

27 Chowdhury, S., Wang, J., Nuccio, S. P., Mao, H. & Di Antonio, M. Short LNA-modified oligonucleotide probes as efficient disruptors of DNA G-quadruplexes. Nucleic Acids Res. 50, 7247–7259 (2022).

28 Heddi, B., Cheong, V. V., Martadinata, H. & Phan, A. T. Insights into G-quadruplex specific recognition by the DEAH-box helicase RHAU: Solution structure of a peptide-quadruplex complex. Proc. Natl. Acad. Sci. U. S. A. 112, 9608–9613 (2015).

29 Keppler, A. et al. A general method for the covalent labeling of fusion proteins with small molecules in vivo. Nat. Biotechnol. 21, 86–89 (2003).

30 Rodriguez, R. et al. A novel small molecule that alters shelterin integrity and triggers a DNA-damage response at telomeres. J. Am. Chem. Soc. 130, 15758–15759 (2008).

31 De Cian, A., Delemos, E., Mergny, J. L., Teulade-Fichou, M. P. & Monchaud, D. Highly efficient G-quadruplex recognition by bisquinolinium compounds. J. Am. Chem. Soc. 129, 1856–1857 (2007).

32 Deng, W., Shi, X., Tjian, R., Lionnet, T. & Singer, R. H. CASFISH: CRISPR/Cas9-mediated in situ labeling of genomic loci in fixed cells. Proc. Natl. Acad. Sci. U. S. A. 112, 11870–11875 (2015).

33 Rosonovski, S., Guneri, D., King, J., Morris, C. J. & Waller, Z. A. E. Affinity-selected peptide ligands specifically bind i-motif DNA and modulate c-Myc gene expression. bioRxiv, 2025.2005.2030.656635 (2025). 10.1101/2025.05.30.656635

34 Qin, G. et al. Targeting specific DNA G-quadruplexes with CRISPR-guided G-quadruplex-binding proteins and ligands. Nat. Cell. Biol. 26, 1212–1224 (2024).

35 Kendrick, S. et al. The dynamic character of the BCL2 promoter i-motif provides a mechanism for modulation of gene expression by compounds that bind selectively to the alternative DNA hairpin structure. J. Am. Chem. Soc. 136, 4161–4171 (2014).

36 Debnath, M. et al. Preferential targeting of i-motifs and G-quadruplexes by small molecules. Chem. Sci. 8, 7448–7456 (2017).

37 King, J. J. et al. DNA G-Quadruplex and i-Motif Structure Formation Is Interdependent in Human Cells. J. Am. Chem. Soc. 142, 20600–20604 (2020).

38 Larson, M. H. et al. CRISPR interference (CRISPRi) for sequence-specific control of gene expression. Nat. Protoc. 8, 2180–2196 (2013).

39 McCarty, N. S., Graham, A. E., Studena, L. & Ledesma-Amaro, R. Multiplexed CRISPR technologies for gene editing and transcriptional regulation. Nat. Commun. 11, 1281 (2020).

40 Prasad, B. et al. A complementary chemical probe approach towards customized studies of G-quadruplex DNA structures in live cells. Chem. Sci. 13, 2347–2354 (2022).

41 Anders, C., Niewoehner, O., Duerst, A. & Jinek, M. Structural basis of PAM-dependent target DNA recognition by the Cas9 endonuclease. Nature 513, 569–573 (2014).

42 Tomasello, G., Armenia, I. & Molla, G. The Protein Imager: a full-featured online molecular viewer interface with server-side HQ-rendering capabilities. Bioinformatics 36, 2909–2911 (2020).

43 Lam, E. Y., Beraldi, D., Tannahill, D. & Balasubramanian, S. G-quadruplex structures are stable and detectable in human genomic DNA. Nat. Commun. 4, 1796 (2013).

44 Peraro, L. et al. Cell Penetration Profiling Using the Chloroalkane Penetration Assay. J. Am. Chem. Soc. 140, 11360–11369 (2018).

45 Gilbert, L. A. et al. Genome-Scale CRISPR-Mediated Control of Gene Repression and Activation. Cell 159, 647–661 (2014).

46 Qi, L. S. et al. Repurposing CRISPR as an RNA-guided platform for sequence-specific control of gene expression. Cell 152, 1173–1183 (2013).

47 Ahmadi, S. E., Rahimi, S., Zarandi, B., Chegeni, R. & Safa, M. MYC: a multipurpose oncogene with prognostic and therapeutic implications in blood malignancies. J. Hematol. Oncol. 14, 121 (2021).

48 Albert, T. et al. The chromatin structure of the dual c-myc promoter P1/P2 is regulated by separate elements. J. Biol. Chem. 276, 20482–20490 (2001).

49 Zhang, X., Spiegel, J., Martinez Cuesta, S., Adhikari, S. & Balasubramanian, S. Chemical profiling of DNA G-quadruplex-interacting proteins in live cells. Nat. Chem. 13, 626–633 (2021).

50 Liberzon, A. et al. The Molecular Signatures Database (MSigDB) hallmark gene set collection. Cell Syst. 1, 417–425 (2015).

51 Pena Martinez, C. D., et al. Human genomic DNA is widely interspersed with i-motif structures. EMBO J. 43, 4786–4804 (2024).

52 Simonsson, T., Pribylova, M. & Vorlickova, M. A nuclease hypersensitive element in the human c-myc promoter adopts several distinct i-tetraplex structures. Biochem. Biophys. Res. Commun. 278, 158–166 (2000).

53 Siddiqui-Jain, A., Grand, C. L., Bearss, D. J. & Hurley, L. H. Direct evidence for a G-quadruplex in a promoter region and its targeting with a small molecule to repress c-MYC transcription. Proc. Natl. Acad. Sci. U. S. A. 99, 11593–11598 (2002).

54 Sun, D. & Hurley, L. H. The importance of negative superhelicity in inducing the formation of G-quadruplex and i-motif structures in the c-Myc promoter: implications for drug targeting and control of gene expression. J. Med. Chem. 52, 2863–2874 (2009).

55 Sutherland, C., Cui, Y., Mao, H. & Hurley, L. H. A Mechanosensor Mechanism Controls the G-Quadruplex/i-Motif Molecular Switch in the MYC Promoter NHE III(1). J. Am. Chem. Soc. 138, 14138–14151 (2016).

56 Hui, W. W. I., Simeone, A., Zyner, K. G., Tannahill, D. & Balasubramanian, S. Single-cell mapping of DNA G-quadruplex structures in human cancer cells. Sci. Rep. 11, 23641 (2021).

57 Pennarun, G. et al. Apoptosis related to telomere instability and cell cycle alterations in human glioma cells treated by new highly selective G-quadruplex ligands. Oncogene 24, 2917–2928 (2005).

58 Yu, Z. et al. Chem-map profiles drug binding to chromatin in cells. Nat. Biotechnol. 41, 1265–1271 (2023).

59 Cuddapah, S. et al. Genomic profiling of HMGN1 reveals an association with chromatin at regulatory regions. Mol. Cell Biol. 31, 700–709 (2011).

60 Kugler, J. E., Deng, T. & Bustin, M. The HMGN family of chromatin-binding proteins: dynamic modulators of epigenetic processes. Biochim. Biophys. Acta 1819, 652–656 (2012).

61 Rosano, D. et al. Long-term Multimodal Recording Reveals Epigenetic Adaptation Routes in Dormant Breast Cancer Cells. Cancer Discov. 14, 866–889 (2024).

62 Berner, A. et al. G4-Ligand-Conjugated Oligonucleotides Mediate Selective Binding and Stabilization of Individual G4 DNA Structures. J. Am. Chem. Soc. 146, 6926–6935 (2024).

63 He, Y. D. et al. Selective Targeting of Guanine-Vacancy-Bearing G-Quadruplexes by G-Quartet Complementation and Stabilization with a Guanine-Peptide Conjugate. J. Am. Chem. Soc. 142, 11394–11403 (2020).

64 Tan, D. J. Y., Das, P., Winnerdy, F. R., Lim, K. W. & Phan, A. T. Guanine anchoring: a strategy for specific targeting of a G-quadruplex using short PNA, LNA and DNA molecules. Chem. Commun. (Camb) 56, 5897–5900 (2020).

65 Zhao, H., Lau, H. L., Zhang, K. & Kwok, C. K. Selective recognition of RNA G-quadruplex in vitro and in cells by L-aptamer-D-oligonucleotide conjugate. Nucleic Acids Res. 52, 13544–13560 (2024).

66 Felsenstein, K. M. et al. Small Molecule Microarrays Enable the Identification of a Selective, Quadruplex-Binding Inhibitor of MYC Expression. ACS Chem. Biol. 11, 139–148 (2016).

67 Flusberg, D. A. et al. Identification of G-Quadruplex-Binding Inhibitors of Myc Expression through Affinity Selection-Mass Spectrometry. SLAS Discov. 24, 142–157 (2019).

68 Robinson, J. et al. G-quadruplex structures regulate long-range transcriptional reprogramming to promote drug resistance in ovarian cancer cells. Genome Biol. 26, 183 (2025).

69 Studier, F. W. Protein production by auto-induction in high density shaking cultures. Protein Expr. Purif. 41, 207–234 (2005).

70 Lingeman, E., Jeans, C. & Corn, J. E. Production of Purified CasRNPs for Efficacious Genome Editing. Curr. Protoc. Mol. Biol. 120, 31 10 31–31 10 19 (2017).

