## Extended Data for "Chemically modified CRISPR-Cas9 enables targeting of individual G-quadruplex and i-motif structures, revealing ligand-dependent transcriptional perturbation"

**Extended Data Table 1. G4-ligands-mediated thermal stabilization of the G4-forming hTelo oligonucleotide *via* FRET-melting.**

$\Delta T_m$ values were determined by FRET-melting assays using 4 µM G4 ligand and 0.4 µM dual-labelled hTelo oligonucleotide. Data represent the mean of three independent experiments (n = 3). Statistical significance was calculated using a two-tailed t-test in GraphPad Prism. p-value: ns > 0.05, * ≤0.05, ** ≤0.01, *** ≤0.001, **** ≤0.0001.

| **G4 Ligand (4 µM)** | **$\Delta T_m$-hTelo ($T_{m4\,\mu M}$ - $T_{m\,0\,\mu M}$)** | **p-value (paired, Two-tailed)** |
|---|---|---|
| PyPDS | 16.48 | *** |
| Cl-PDS$_2$ | N/A | * |
| Cl-PDS$_4$ | 17 | * |
| PhenDC3 | 35.01 | ** |
| Cl-PhenDC3$_2$ | 23.43 | ** |
| Cl-PhenDC3$_4$ | 28.93 | **** |

**Extended Data Table 2 G4-ligands-mediated thermal stabilization of the G4-forming BCL2 oligonucleotide *via* FRET-melting.**

$\Delta T_m$ values were determined by FRET-melting assays using 4 µM G4 ligand and 0.4 µM dual-labelled BCL2 oligonucleotide. Data represent the mean of three independent experiments (n = 3). Statistical significance was calculated using a two-tailed t-test in GraphPad Prism. p-value: ns > 0.05, * ≤0.05, ** ≤0.01, *** ≤0.001, **** ≤0.0001.

| **G4 Ligand (4 µM)** | **$\Delta T_m$-BCL2 ($T_{m4\,\mu M}$ - $T_{m\,0\,\mu M}$)** | **p-value (paired, Two-tailed)** |
|---|---|---|
| PyPDS | 4.66 | * |
| Cl-PDS$_2$ | 4.47 | ns |
| Cl-PDS$_4$ | 5.08 | * |
| PhenDC3 | 30.64 | ** |

| | | |
|---|---|---|
| Cl-PhenDC3$_2$ | 25.58 | ** |
| Cl-PhenDC3$_4$ | 29.24 | ** |

**Extended Data Table 3 G4-ligands-mediated thermal stabilization of the G4-forming c-KIT2 oligonucleotide *via* FRET-melting.**

$\Delta T_m$ values were determined by FRET-melting assays using 4 µM G4 ligand and 0.4 µM dual-labelled c-KIT2 oligonucleotide. Data represent the mean of three independent experiments (n = 3). Statistical significance was calculated using a two-tailed t-test in GraphPad Prism. p-value: ns > 0.05, * ≤0.05, ** ≤0.01, *** ≤0.001, **** ≤0.0001.

| G4 Ligand (4 µM) | $\Delta T_m$ c-KIT2 ($T_{m4\,\mu M}$ - $T_{m\,0\,\mu M}$) | p-value (paired, Two-tailed) |
|---|---|---|
| PyPDS | 14.93 | ** |
| Cl-PDS$_2$ | 16.4 | ** |
| Cl-PDS$_4$ | 11.18 | ** |
| PhenDC3 | 20.91 | *** |
| Cl-PhenDC3$_2$ | 13.71 | ** |
| Cl-PhenDC3$_4$ | 12.37 | ** |

**Extended Data Table 4 G4-ligands-mediated thermal stabilization of the G4-forming c-MYC-Pu22 oligonucleotide *via* CD-melting.**

$\Delta T_m$ values were determined by CD-melting assays using increasing concentration (µM) of G4 ligand and 2 µM of c-MYC(Pu22) oligonucleotide. Data represent the mean of three independent experiments (n = 3). Statistical significance was calculated using a two-tailed t-test with Welch test correction in GraphPad Prism. p-value: ns > 0.05, * ≤0.05, ** ≤0.01, *** ≤0.001, **** ≤0.0001.

| | Concentration (µM) | | | | | | | |
|---|---|---|---|---|---|---|---|---|
| | 0.5 | | 1.0 | | 2.0 | | 4.0 | |
| G4 Ligand | $\Delta T_m$ | p-value | $\Delta T_m$ | p-value | $\Delta T_m$ | p-value | $\Delta T_m$ | p-value |
| PyPDS | 6.4 | *** | 12.3 | **** | 14.4 | **** | 17.6 | **** |
| Cl-PDS$_2$ | 4.6 | *** | 8.8 | **** | 11.2 | **** | 15.9 | **** |
| Cl-PDS$_4$ | -0.3 | ns | 7.3 | **** | 10.4 | **** | 13.1 | **** |

| | | | | | | | | |
|---|---|---|---|---|---|---|---|---|
| **PhenDC3** | 11.0 | ** | 41.3 | **** | 59.5 | ns (not accurate, no melting Tm>95 °C) | n/a | n/a |
| **Cl-PhenDC3$_2$** | 9.6 | ** | 39.5 | **** | 57.1 | ns (not accurate, no melting Tm>95 °C) | n/a | n/a |
| **Cl-PhenDC3$_4$** | 6.0 | ** | 37.3 | **** | 44.8 | ns (not accurate, no melting Tm>95 °C) | n/a | n/a |

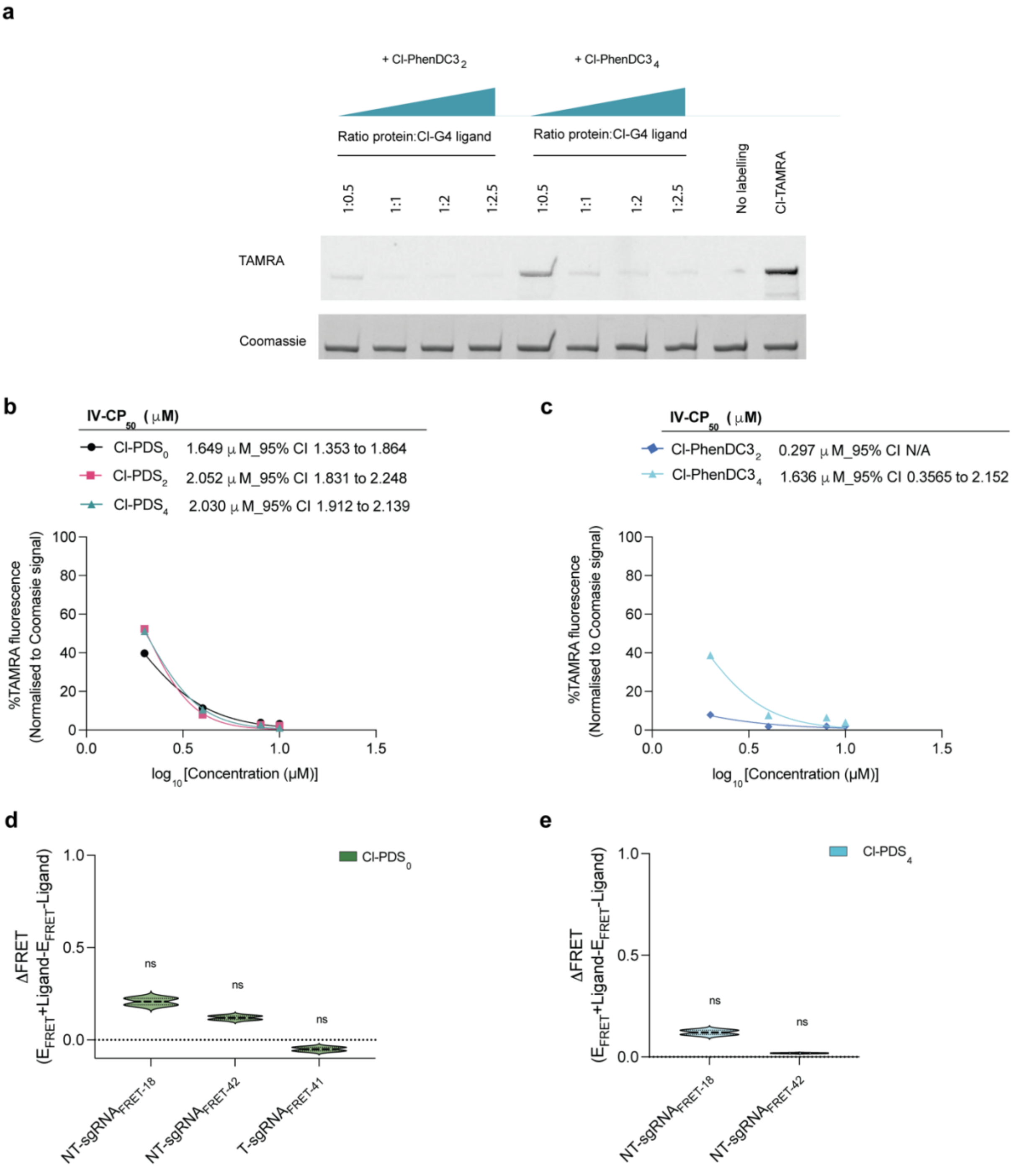


**Extended data Figure 1 ATENA biochemical characterization a,** SDS-Page gel of the Cl-PhenDC3$_n$ competition assay showing each sample's fluorescent level acquired in the TAMRA channel (542 nm) and the corresponding protein level (Coomassie staining). (n=2) **b, c,** Plot of the band intensity value in the TAMRA channel indicative of dCas9-Halo labeling efficiency *in vitro* for Cl-PDS$_2$ and Cl-PhenDC3$_2$, respectively, and relative IV-CP$_{50}$. Each band intensity was quantified using Image Studio software and normalized for the corresponding Coomassie signal, expressed as a percentage (with 100% labeling corresponding to the positive control). IV-CP$_{50}$ values were determined using nonlinear regression (dose-response inhibition curves with constrained fitting) in GraphPad Prism (n = 2) with $R^2$

values of 0.9957 for Cl-PDS$_0$, $R^2$ values of 0.9956 for Cl-PDS$_2$ and $R^2$ values of 0.9988 for Cl-PDS$_4$. $R^2$ values N/A for Cl-PhenDC3$_2$ and 0.9582 for Cl-PhenDC3$_4$. **d, e** ΔFRET efficiency of the decorated dCas9-PDSx(with Cl-PDS$_0$ and Cl-PDS$_4$, respectively) targeting c-KIT2-G4. The values indicated were extrapolated from the band intensity measured in the Cy3 and Cy5 channels (Typhoon FLA 9500). The signals in both channels were normalised for the background and the sgRNA NTC control. The normalised fluorescence values were then used to calculate the ΔFRET efficiency as follows for each sgRNA: FRET-Efficiency(E)+ligand$_{sgRNAx}$ – FRET-Efficiency(E)-ligand$_{sgRNAx}$ (n=2). The data presented are the mean of n = number of independent experiments. Statistical significance was calculated using a Welch-corrected two-tailed t-test in GraphPad Prism; p-value: ns > 0.05, * ≤0.05, ** ≤0.01, *** ≤0.001, **** ≤0.0001.

.

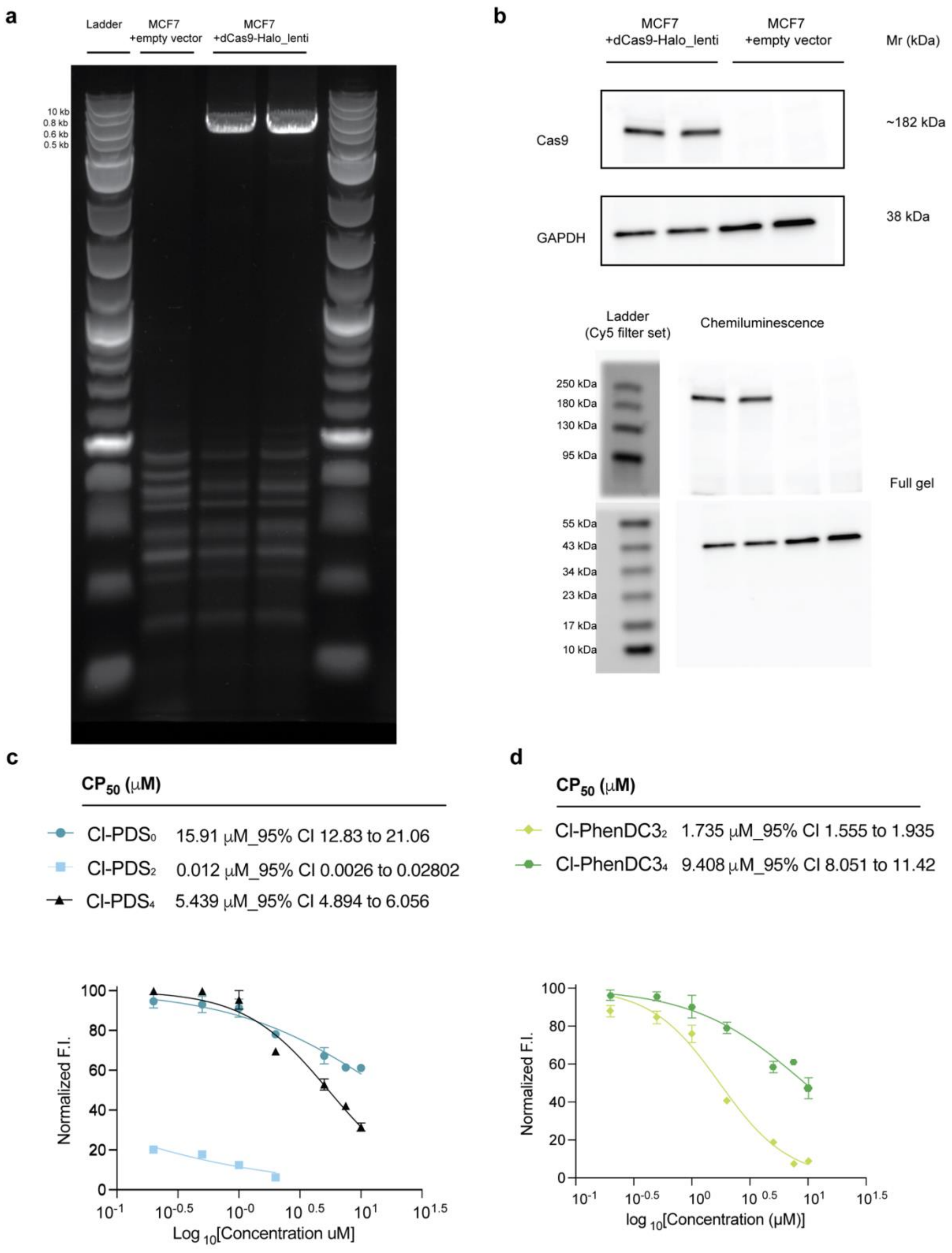

a
Ladder
MCF7 +empty vector
MCF7 +dCas9-Halo_lenti
10 kb
0.8 kb
0.6 kb
0.5 kb
b
MCF7 +dCas9-Halo_lenti
MCF7 +empty vector
Mr (kDa)
Cas9
~182 kDa
GAPDH
38 kDa
Ladder (Cy5 filter set)
Chemiluminescence
250 kDa
180 kDa
130 kDa
95 kDa
55 kDa
43 kDa
34 kDa
23 kDa
17 kDa
10 kDa
Full gel
c
$CP_{50}$ (μM)
Cl-PDS$_0$ 15.91 μM_95% CI 12.83 to 21.06
Cl-PDS$_2$ 0.012 μM_95% CI 0.0026 to 0.02802
Cl-PDS$_4$ 5.439 μM_95% CI 4.894 to 6.056
Normalized F.I.
Log $_{10}$[Concentration uM]
d
$CP_{50}$ (μM)
Cl-PhenDC3$_2$ 1.735 μM_95% CI 1.555 to 1.935
Cl-PhenDC3$_4$ 9.408 μM_95% CI 8.051 to 11.42
Normalized F.I.
log $_{10}$[Concentration (μM)]

**Extended data Figure 2 ATENA optimization in mammalian cells. a,** Genotyping of MC7 cells transduced with either empty vector or dCas9-Halo lentiviral construct showing the amplified expected band at 5.1 kb (SC_dCas9genotyping.Fw, SC_dCas9genotyping.Rev) in case of successful integration. **b,** Western blot of either MCF7+dCas9-Halo stable cell line or MCF7+empty vector to confirm protein expression of dCas9-Halo. Gel images were acquired using Image Quant LAS 4000 (Cytiva) following the methodology described in the Methods section. **c, d,** Labeling efficiency of dCas9–Halo in live cells treated with Cl-PDS$_n$ or Cl-PhenDC3$_n$ and the corresponding CP$_{50}$ values. Cells labeled with the Cl-OG fluorophore were analyzed by flow cytometry (see Methods), and data were processed in FlowJo. Mean fluorescence values from two biological replicates (each with three technical replicates) were normalized to the positive-control signal and expressed as percent labeling. CP$_{50}$ values were obtained by nonlinear regression (dose–response inhibition curves with constrained fitting) in GraphPad Prism (n = 2). The fits yielded $R^2$ values of 0.9374 for Cl-PDS$_0$, 0.8598 for Cl-PDS$_2$, 0.9686 for Cl-PDS$_4$, 0.9775 for Cl-PhenDC3$_2$, and 0.9406 for Cl-PhenDC3$_4$. Data points at concentrations $\geq 5$ µM were excluded from the Cl-PDS$_2$ fit because the compound was cytotoxic at those levels, as confirmed by its IC$_{50}$ values calculated (see SI table S7).

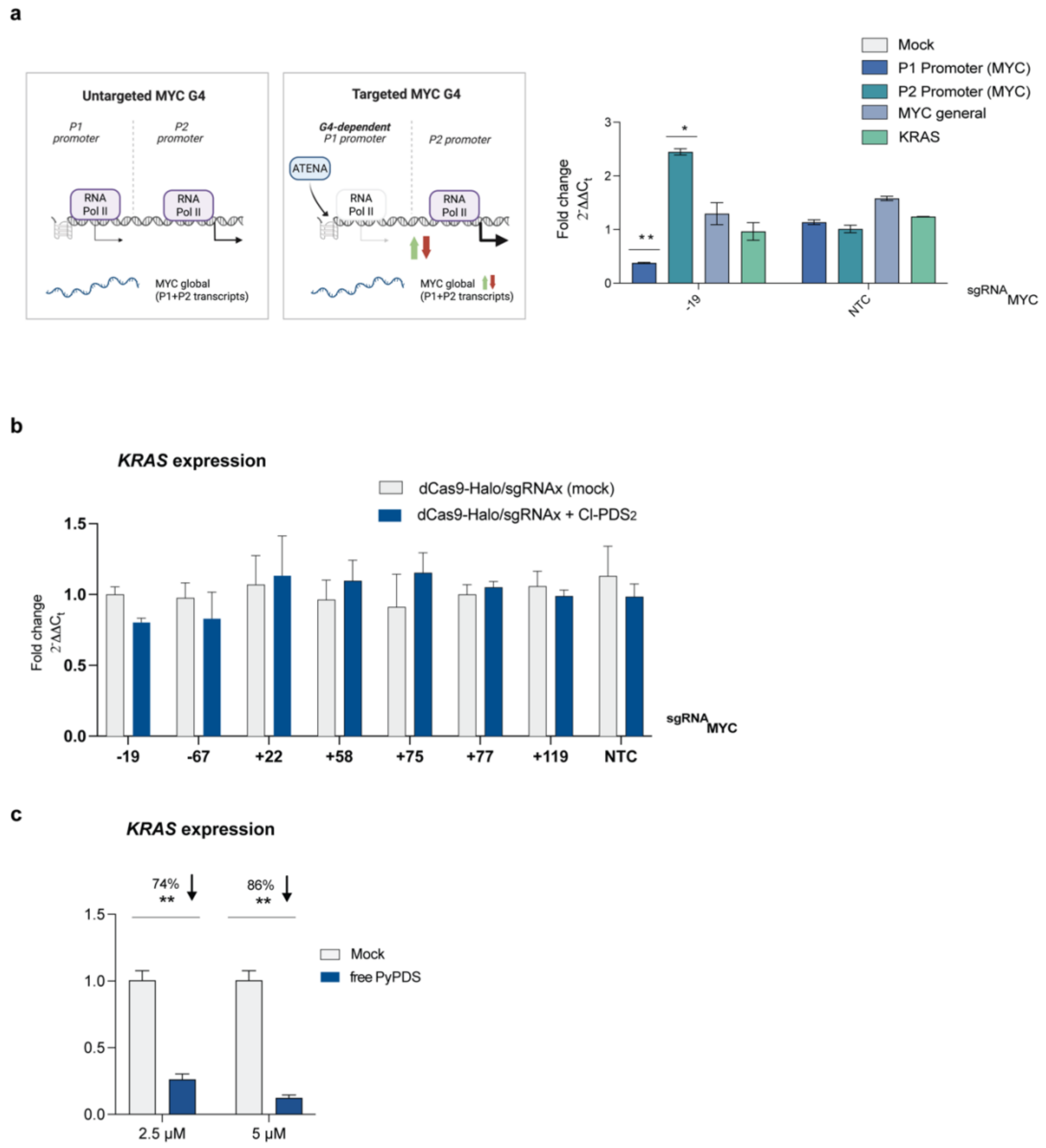


**Extended data Figure 3 P2-compensation and limited off-target activity following ATENA-mediated MYC-G4 targeting. a,** (left) proposed model in which P2-driven transcription compensates for the reduction of P1-mediated transcription that follows MYC-G4 targeting. (right) RT-qPCR of the indicated genes in MCF7 cells stably expressing dCas9-Halo transfected with $sgRNA_{MYC-19}$ or sgRNA NTC and incubated for 48h in the presence of 2.5 µM of Cl-$PDS_2$ or DMSO (mock). The expression values are represented as fold change ($2^{-\Delta\Delta Ct}$) with respect to the mock (DMSO-treated) transfected samples and after normalization for the housekeeping gene *(GAPDH*). n=2, biological replicates, each with three technical replicates. **b,** RT-qPCR for *KRAS* expression in MCF7 cells stably expressing dCas9-Halo transfected with the indicated sgRNAs and incubated for 48h in the presence of (2.5 µM) Cl-$PDS_2$. The expression values are represented as fold change ($2^{-\Delta\Delta Ct}$) with respect to the mock (DMSO-treated) and normalized for the housekeeping gene (*GAPDH).* n=3, biological replicates, each with three technical replicates.**c,** RT-qPCR for *KRAS* expression in MCF7 cells incubated for 24h in the presence of (2.5 µM) free PyPDS. The expression values are represented as fold change ($2^{-\Delta\Delta Ct}$) with respect to the mock (DMSO-treated) and after normalization for the housekeeping gene *(GAPDH*). n=3, biological replicates, each with two technical replicates. The data presented are the mean of n = number of independent biological samples. Statistical significance was calculated using a Welch-corrected two-tailed t-test in GraphPad Prism; p-value: ns > 0.05, * ≤0.05, ** ≤0.01, *** ≤0.001, **** ≤0.0001.

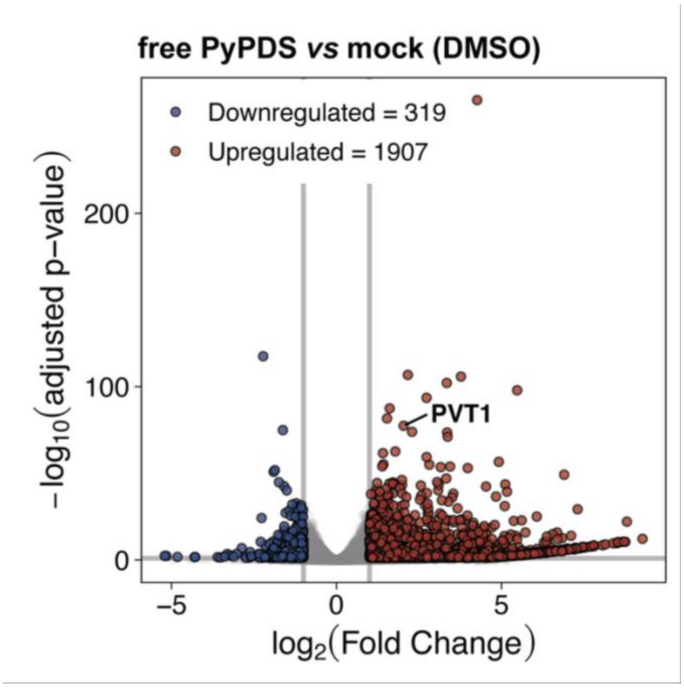


**Extended data Figure 4 Investigating free PyPDS effect on global mRNA expression in MCF7 cells**. Volcano plot showing the number of genes differently expressed upon treatment of MCF7 cells with (2.5 uM) free PyPDS. The plot was generated by comparing Cl-$PDS_2$-treated samples and DMSO-treated (mock) samples using DESeq2 with FDR=0.05. Highlighted the *PVT1* gene, whose expression was previously shown to be upregulated after PDS treatment [35].

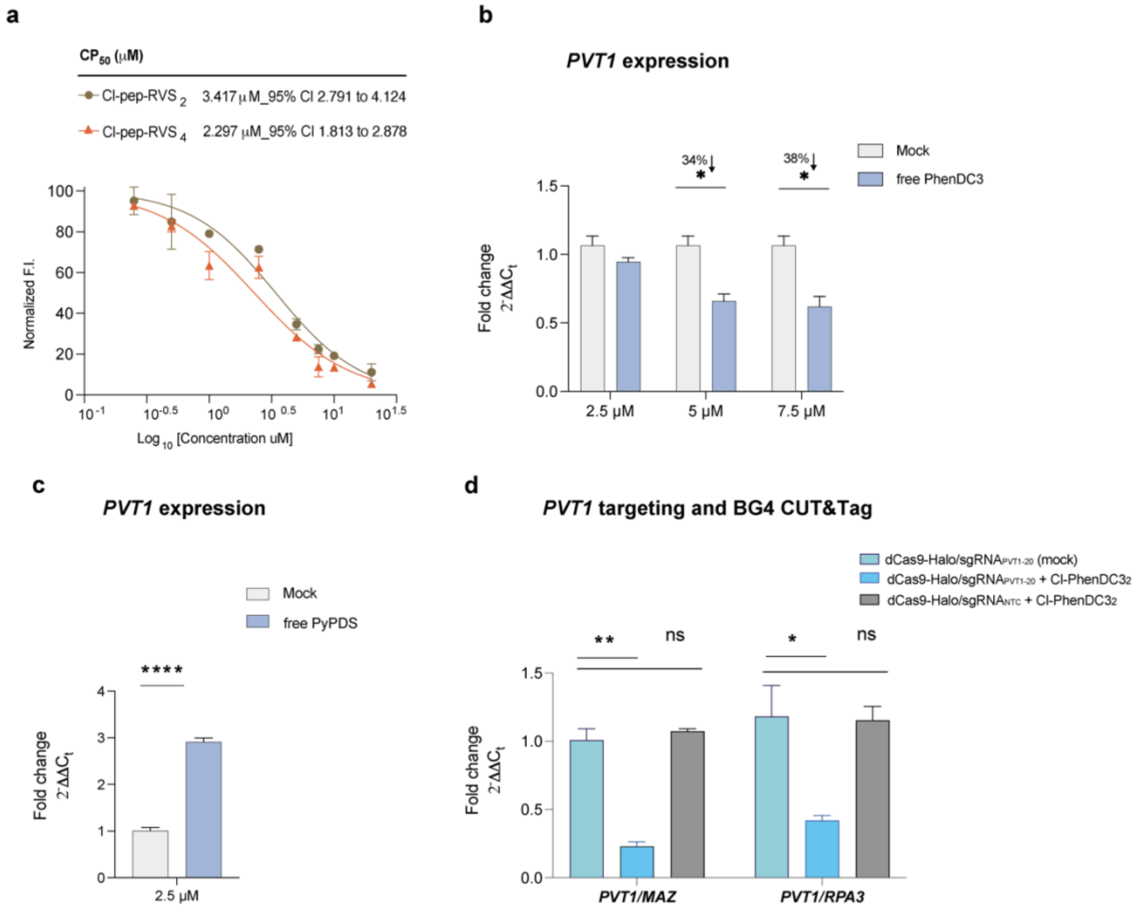


**Extended data Figure 5 *ATENA* targets the *c-Myc* i-motif and *lncPVT1*. a,** Labeling efficiency of dCas9–Halo in live cells treated with Cl-pep-RVS$_2$ or Cl-pep-RVS$_4$ and the corresponding CP$_{50}$ values. Cells labeled with the Cl-OG fluorophore were analyzed by flow cytometry (see Methods), and data were processed in FlowJo. Mean fluorescence values from two biological replicates (each with three technical replicates) were normalized to the positive-control signal and expressed as percent labeling. CP$_{50}$ values were obtained by nonlinear regression (dose–response inhibition curves with constrained fitting) in GraphPad Prism (n = 2). The fits yielded $R^2$ values of 0.9548 for Cl-pep-RVS$_2$ and 0.9491 for Cl-pep-RVS$_4$. **b,** RT-qPCR for *PVT1* expression in MCF7 cells treated with increasing concentration of free PhenDC3 for 24h or DMSO (mock). The expression values are represented as fold change ($2^{-\Delta\Delta Ct}$) with respect to the mock (DMSO-treated) transfected samples and after normalization for the housekeeping gene *(GAPDH)*. n=2, biological replicates, each with two technical replicates. **c**, RT-qPCR for *PVT1* expression in MCF7 cells treated with (2.5 µM) free PyPDS for 24h or DMSO (mock). The expression values are represented as fold change ($2^{-\Delta\Delta Ct}$) with respect to the mock (DMSO-treated) transfected samples and after normalization for the housekeeping gene *(GAPDH)*. n=2, biological replicates, each with two technical replicates. **d,** BG4 CUT&Tag-qPCR for MCF7 cells stably expressing dCas9-Halo transfected with either sgRNA$_{PVT1-20}$ or sgRNA NTC and treated with DMSO (mock) or (2.5 µM) Cl-PhenDC3$_2$. BG4 accessibility was analyzed for *PVT1* and normalized to two G4s in control gene sites (*MAZ* and *RPA3*). n=2, biological replicates each with three technical replicates for BG4 and one for the negative (no BG4 treatment). Data presented are mean of n = number of independent biological samples. Statistical significance was calculated using a Welch-corrected two-tailed t-test in GraphPad Prism; p-value: ns > 0.05, * ≤0.05, ** ≤0.01, *** ≤0.001, **** ≤0.0001.

a

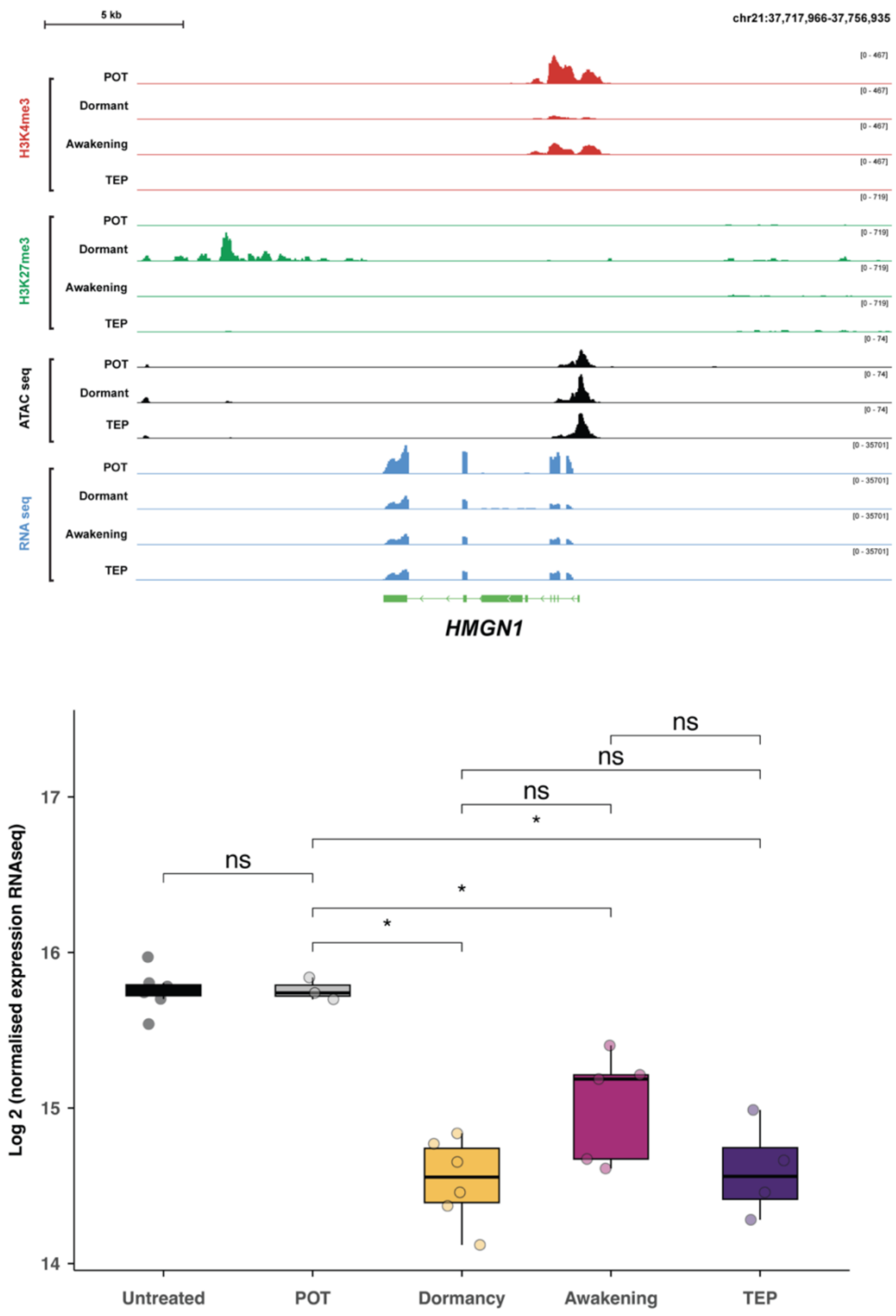


**Extended data Figure 6 Epigenetic and transcriptional landscape at the *HMGN1* locus in MCF7 cells under different Estrogen-deprivation conditions. a,** Genome browser tracks display group-auto-scaled CUT&Tag signal intensity for the histone mark H3K4me3 (red), and H3K27me3 (green); ATAC-seq signal intensity (black) and RNA-seq coverage (blue) across the *HMGN1* locus. Each track corresponds to a distinct time point in Estrogen-deprivation treatment (POT (Day 0), Dormancy (Day 43), Awakening, and TEP - Terminal End Point, as indicated on the left. Peaks in the H3K4me3 and RNA-seq tracks are associated with active promoters and transcriptional activity, respectively, while signal in the H3K27me3 track indicates repressive chromatin. Gene models are shown at the bottom, with exon-intron structure and transcriptional direction indicated. **b,** Boxplots showing gene expression levels of *HMGN1* gene as log-transformed DESeq2 and normalized counts from bulk RNAseq in MCF7 cells subjected to Estrogen deprivation treatment. Untreated: MCF7 grown in complete media with Estradiol, POT: Starting population of MCF7 grown in complete media with Estradiol. Dormancy: MCF7 cells were grown in estrogen-deprived medium for 1-3 months. Awakening: Dormant populations that resumed proliferation. TEP (Terminal End Points): awakening populations subcultured for at least 30 days. Statistical significance was determined using the Wilcoxon rank-sum test, with p-values indicated by asterisks (*).
